# Analyzing Assay Specificity in Metabolomics using Unique Ion Signature Simulations

**DOI:** 10.1101/2021.03.19.434579

**Authors:** Premy Shanthamoorthy, Adamo Young, Hannes Röst

## Abstract

Targeted, untargeted and data-independent acquisition (DIA) metabolomics workflows are often hampered by ambiguous identification based on either MS1 information alone or relatively few MS2 fragment ions. While DIA methods have been popularized in proteomics, it is less clear whether they are suitable for metabolomics workflows due to their large precursor isolation windows and complex co-isolation patterns. Here, we quantitatively investigate the conditions necessary for unique metabolite detection in complex backgrounds using precursor and fragment ion mass-to-charge separation, comparing three benchmarked MS methods (MS1, MRM, DIA). Our simulations show that DIA outperformed MS1-only and MRM-based methods with regards to specificity by a factor of ~2.8-fold and ~1.8-fold, respectively. Additionally, we show that our results are not dependent on the number of transitions used or the complexity of the background matrix. Finally, we show that collision energy is an important factor in unambiguous detection and that a single collision energy setting per compound cannot achieve optimal pairwise differentiation of compounds. Our analysis demonstrates the power of using both high resolution precursor and high resolution fragment ion *m/z* for unambiguous compound detection. This work also establishes DIA as an emerging MS acquisition method with high selectivity for metabolomics, outperforming both DDA and MRM with regards to unique compound identification potential.

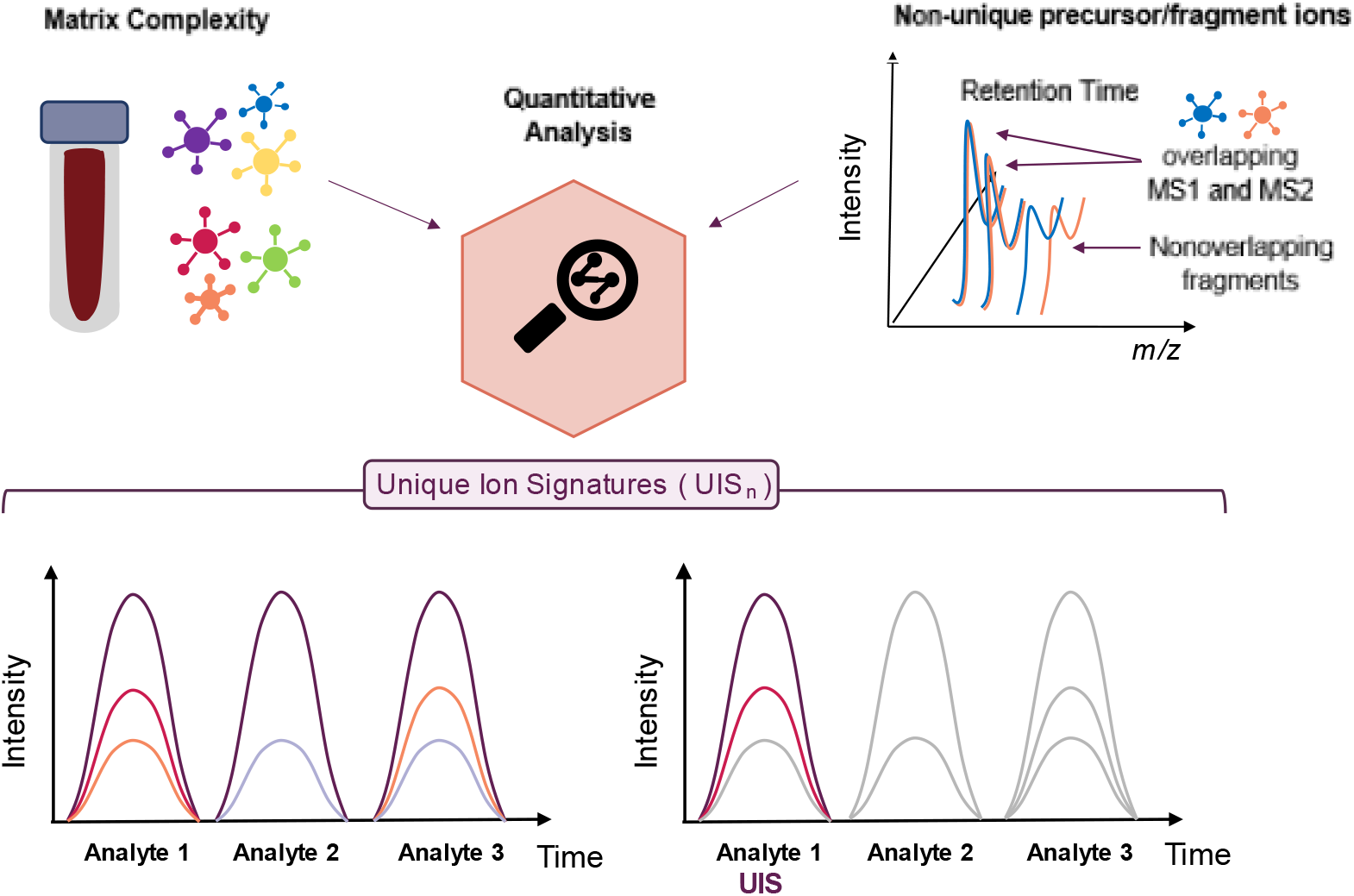

## Introduction

Liquid chromatography coupled to tandem mass spectrometry (LC-MS/MS) allows for the robust analysis of complex samples in metabolomics. LC/MS-based metabolomics allows researchers to explore a large fraction of chemical diversity, uncovering fundamental metabolism, regulation, genetics, and interspecies analyte transfer in complex systems of global importance, such as nutrient cycling, wastewater treatment and the human microbiome.^1–2^

However, the amazing diversity of the metabolome and its lack of a generic polymer template, as in the genome or proteome, greatly complicates the confident identification of unique metabolites required to provide meaningful biological data. Untargeted workflows focused on the precursor ion (MS1-based) are comprehensive, but often unspecific with compound detection hampered by their reliance on a single precursor *m/z* value. An increase in confidence occurs with the addition of fragment ion (MS2) data (an MS2 match in literature, towards a level 2 identification with orthogonal information such as retention time or collision cross-section values).^3^ However, MS2 data acquired using data-dependent workflows (DDA) heavily relies on stochastic MS1 data collection, with a high degree of variance based on sample complexity. On the other hand, although targeted methods (multiple/parallel reaction monitoring; MRM/PRM) using MS2 data are highly specific, they are strongly limited in both mass accuracy and analyte throughput for traditional MRM with a focus on few fragments, generally unsuitable for omics type of analysis.

Data-independent acquisition (DIA) is a next-generation MS method with capabilities of capturing the complete precursor and fragment ion signal (MS1 and MS2) in a single run. It uses a set of pre-programmed isolation windows which span the whole mass range, thus ensuring that any precursor is fragmented. In each cycle, a set of high resolution fragment ion spectra is acquired for every isolation window, and the process is repeated once the last isolation window has been acquired, generating a series of fragment ion spectra with the same precursor isolation window. This method allows high analyte throughput with reproducible and consistent quantification to facilitate metabolite discovery.^4–10^

Current DIA methods for metabolomics aim to reduce interference in the overlapping *m/z* and RT space. One all-ion fragmentation MS method named MS^E^ involves alternating scans at low or high collision energy for simultaneous precursor and fragment ion data acquisition within a single analytical run.^11^ MS^E^ maximizes data collection efficiency and duty cycle by taking advantage of the characteristic high acquisition speed and mass accuracy of a quadrupole time-of-flight (Q-TOF) instrument, making it suitable for a large number of highly complex samples.^10–11^ However, as fragmentation is performed for all precursor ions within a wide *m/z* window, the acquired nonselective and highly complex fragment ion spectra may result in poor specificity, which then requires efficient post-acquisition data processing software tools.^5^ An alternative technique to obtain DIA-MS/MS spectra is called SWATH-MS. Initially applied in the proteomics field, this method commonly uses 25 Da consecutive precursor ion isolation windows with the targeted extraction of precursors (25 ppm), and allows for accurate fragment-ion based targeted analysis in a high-throughput, unbiased manner. The associated analysis software OpenSWATH is capable of targeted precursor and fragment ion extraction from DIA data, and subsequent probabilistic scoring of the resulting peaks.^12^ In comparison to MS^E^, SWATH uses narrower MS1 mass isolation windows, maintaining the ability to obtain MS2 data from a wide precursor mass range. Although data is still highly multiplexed, this workflow has been effectively used in metabolomics with increased versatility.^13^ In our work, our interpretation of DIA is of targeted SWATH-like methods.

MS1-only, MRM/PRM and DIA methods all use a combination of evidence based on precursor ion signals, fragment ion signals or a combination of the two for compound detection. Here, we will investigate the required conditions to uniquely detect a compound among a given set of expected compounds in a complex sample using precursor and fragment ion signals. While relying on multiple characteristic fragment ions is a generally accepted technique to increase certainty in targeted metabolomics assay generation, it is unclear how many transitions are “enough” and few studies have tried to quantify the effect of the acquisition method on compound detection on a large scale.^14–16^

Sherman *et al.* first introduced the concept of using information content as criterion to select suitable transitions (the combined representation of a compound with both MS1 and MS2 data) for targeted methods in proteomics.^14–16^ This work is based on the concept of unique ion signatures (UIS), referring to the combinations of ions that map uniquely to one analyte (a peptide in their work), for a given analyte background. In both cases, the selection of assays with minimal interference with other analytes averted the high likelihood of ambiguous detection due to multiple analytes sharing a particular combination of transitions. Here, we introduce the UIS concept for metabolomics, and use it to calculate non-redundant theoretical assays for thousands of compounds in a given metabolomic background. Specifically, we quantify the capabilities of metabolite detection using mass accuracy to compare current methods in metabolomics. The exclusion of interfering candidates during identification facilitates increased confidence, towards Level 1 or Level 0 identifications with the use of internal standards or pure isolation respectively.^3^ These analyses assess the specificity and the power to detect analytes with MS1-only, or with MS2, in addition to assessing the potential of novel combinatorial approaches (DIA) in the field of metabolomics. Methods, such as DDA, where data acquisition is dependent on precursor abundance and sampling biases, are not included in our analyses.

## Experimental Section

### UIS_*n*_

Using the NIST 17 LC/MS library (~14,000 compounds, 600,000 spectra), compounds and spectra were filtered to retain only structurally different compounds measured in positive-ion mode on a high resolution instrument type (higher energy dissociation - HCD and Q-TOF) and a collision energy of 35eV+/−5eV to obtain a “background metabolome” of 10186 compounds (21334 unique precursors, including in-source fragments).^17^ After filtering for common adduct types (H+, Na+), 9156 queries (individual metabolites in the given metabolome) were used for simulations by setting realistic mass tolerances associated with different MS methods (Fig. 1A-B). Fragment ion spectra were additionally filtered to only include valid transitions greater than 10% of the maximum relative intensity in the fragment spectrum, and the unfragmented precursor ion signal was removed. Precursor ions derived from in-source fragments were included in the background metabolome to assess the effect of in-source fragmentation on unique detection (as defined by NIST 17).^17^

**Figure 1.**
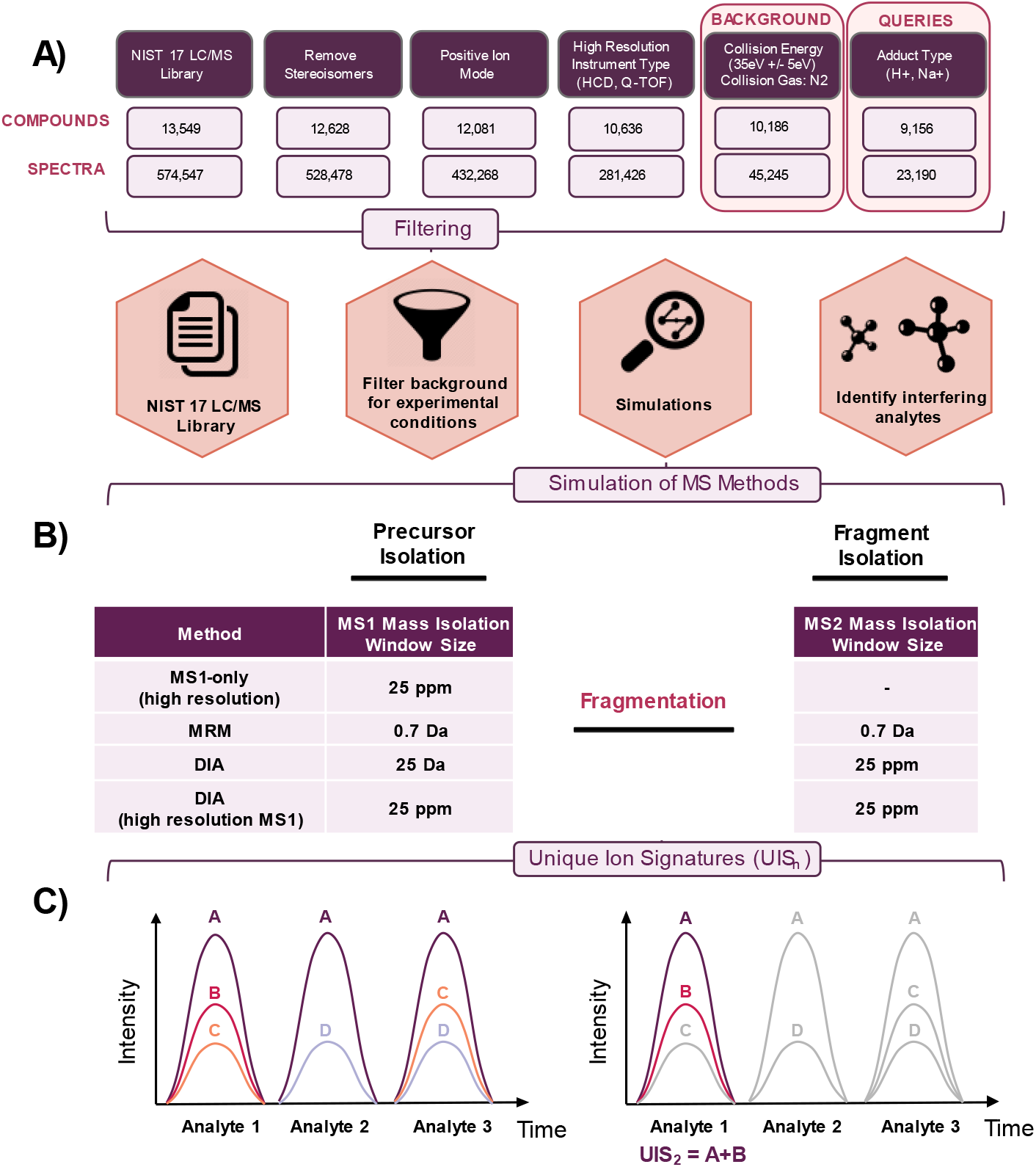
Unique Ion Signature Analysis Pipeline. **(A)** Using the NIST 17 LC/MS library (~14,000 compounds, 600,000 spectra), compounds and spectra were filtered for the removal of stereoisomers, and experimental settings of positive-ion mode, high resolution instrument type (HCD and Q-TOF) and collision energy of 35eV+/−5eV for a background of 10186 compounds. **(B)** After isolating for common adduct types (H+, Na+), 9156 queries, individual metabolites in the given metabolome, were used for simulations by setting realistic mass tolerances associated with different MS methods (Table S-1, MRM – Multiple Reaction Monitoring, DIA – Data Independent Acquisition). **(C)** The UIS concept was used, here exemplified using three analytes that have some transitions in common. UISn is defined as a set of top *n* transitions that map exclusively to one analyte in the metabolome to be analyzed. Assuming a metabolome consisting of three analytes that resolve on the chromatography, using the top *n* transitions - Analyte 1 has no UIS1, a UIS2 of A and B, and a UIS3 of A-C. Note that transition A is not a unique ion signature because this signal can be explained by either Analyte 1, Analyte 2 or Analyte 3. *Figure adapted from:* Röst, H.; Malmström, L.; Aerbersold, R*. Mol. Cell. Proteom.* **2012**.

Each query was independently compared against the full 10186 compounds in the background metabolome using the UIS concept for the tested MS methods (MS1-only, MRM, DIA). UIS_n_ is defined as a set of top *n* analyte transitions (precursor and fragment *m/z*) that map exclusively to one metabolite in the metabolome to be analyzed within the constraints of a given mass resolution.^14–15^ Appropriate values for mass accuracy were used for both the precursor *m/z* window (MS1) and the fragment *m/z* window (MS2) based on the resolution of commercially available instrumentation (triple quadrupole - QQQ or Orbitrap/QTOF). For QQQ instruments we chose an isolation width of 0.7 Da and for high resolution instruments we conservatively assumed a resolution achievable by all high resolution instruments of 40 000 and a corresponding extracted ion chromatogram width of 25 ppm. To simulate instruments with higher resolution, we also explored extraction windows up to 1 ppm (Fig. S-4, Table S-3). However, while mass accuracy is higher than 25 ppm, for targeted extraction the full peak is required for an extracted ion chromatogram (XIC), which is why an extraction window based on resolution and not mass accuracy was used. Additionally, when using a high resolution MS2 (1 ppm), we show that the difference between using an MS1 signal of 25 ppm and 1 ppm is less than 1% (Fig. S-4, Table S-3). Isolation window sizes were set accordingly, centered at the corresponding precursor and fragment *m/z* values of the query, with the number of transitions defined by a value of *n* (UIS_n_). The following MS1/MS2 mass isolation windows were set (in daltons - Da, or parts per million of 1 Da - ppm): MS1-only: 25 ppm/-; Multiple Reaction Monitoring (MRM): 0.7 Da/0.7 Da; and Data-Independent Acquisition (DIA): 25 Da/25 ppm, 25 ppm/25 ppm (Table S-1).

Simulations provided a measure of uniqueness for each metabolite based on the number of compounds found in the background that were not differentiable from the query at the given parameters. For example, in Fig. 1C, assuming a metabolome consists of three analytes, our simulation would determine that for analyte 1, the transition pair A and B (using the two most abundant transitions) produces a UIS_2_, as no other compound contains the combination of these two transitions. With no interfering compounds, the UIS_2_ A-B allows for the unique detection of analyte 1 (Fig. S-12). With this method, the number of hits in the background and the percentage of compounds in the NIST library (queries) with no interference (the background for each query based on the given parameters), was determined for each method. Analyses and visualization discussed herein were performed using Python 3.6. All scripts are available through Github [BSD License 3.0] at: https://github.com/premyshan/DIAColliderMetabo.

### Theoretical Saturation

#### Number of Transitions

Simulations were performed with respect to the number of transitions defined for UIS_n_ using different acquisition methods that use MS2 information (MRM, DIA). Fragment spectra for each query were filtered to include the top *n* transitions greater than 10% of the maximum relative intensity, ranging from *n*=1-8 for UIS_n_. Simulations were then conducted using these individual queries to determine corresponding background metabolites that interfere within the set MS1/MS2 isolation windows for MRM (0.7 Da/0.7 Da) or DIA (25 Da/25 ppm, 25 ppm/25 ppm). The percentage of unique compounds was then calculated (100% = 9156 queries).

#### Matrix Complexity

UIS simulations were conducted to determine the performance of different MS methods (measured by the percentage of unique compounds) in relation to matrix complexity, as indicated by the number of compounds included in the background. The following methods were used to perform 100 simulations for each subset of the NIST 17 LC/MS library as background (ranging from 1000-9000 compounds) - MS1-only: 25 ppm/-; MRM: 0.7 Da/0.7 Da; and DIA: 25 Da/25 ppm, 25 ppm/25 ppm. For methods using MS2, three transitions were used (UIS_3_). The median percentage of unique compounds was calculated for each method (for all subsets), and saturation effects were then modelled using statsmodels (Python module) with a logarithmic transformation (with extrapolation to 15,000 compounds).

#### Collision Energy

Compounds from the NIST 17 LC/MS library were filtered by experimental conditions (positive ion mode and removal of stereoisomers), mass isolation window size (MS1 = 25 Da), adduct (H+) and acquisition instrument (Q-TOF) to calculate optimal collision energies (CE) for each compound (Fig. 5A, Fig. S-8). Individual compounds were compared against each of their interfering compounds using two values to create a similarity matrix - (i) the negative absolute difference in CE and (ii) their cosine similarity score based on their spectra. First, comparisons with a minimum absolute difference in CE were chosen for each row and column (below a maximum CE difference threshold, Fig. S-8). From these comparisons, pairwise-optimal CE (POCE) were determined based on the overall minimum similarity score.

#### Results and Discussion

To investigate the problems of assay redundancy and specificity in metabolomics, we used computational models to calculate nonredundant theoretical assays using the UIS concept for a given metabolomic background. For this analysis, we simulated different MS methods (MS1-only, MRM, DIA) using the NIST 17 LC-MS library as a background (10186 compounds at collision energy = 35eV+/−5eV), using realistic values for both the precursor *m/z* window (MS1) and the fragment *m/z* window (MS2), measured in daltons (Da), or parts per million of a dalton (ppm) (Fig. 1). In our simulations, we selected single query molecules to be tested against a complex background (e.g. the full NIST 17 library).

We compared the selected precursor and fragment ion coordinates of the query compound against each compound in the background set (excluding stereoisomers which would not be distinguishable by MS). We report any overlap in precursor and fragment ion coordinates at a given *m/z* threshold, which can occur at a level of MS2 only, with separability by high resolution MS1 data (Fig. 2A), or at both levels of MS1 and MS2 (Fig. 2B).

**Figure 2.**
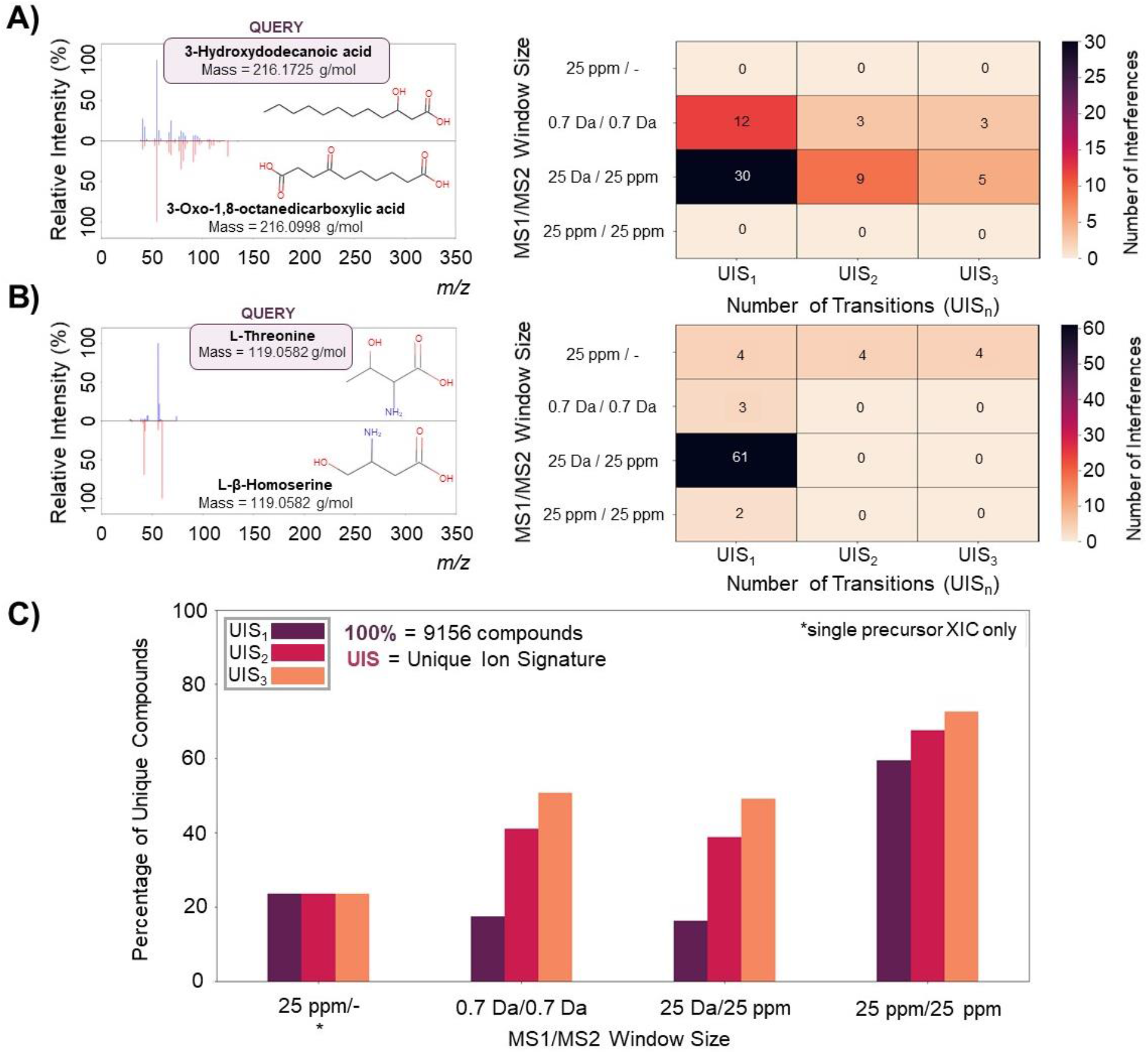
MS1 and MS2 contribute orthogonally to unambiguous compound detection. Pairs of queries and their interfering analytes are displayed with corresponding MS2 maps, with relative intensity (as a percentage of the most abundant ion) on the y-axis and fragment *m/z* on the x-axis. The number of interfering compounds from UIS simulations are represented by the gradient from orange to purple signifying low to high interference for varying MS methods using MS1 and/or MS2 information (MS1-only, MRM, DIA). **(A)**3-Hydroxydodecanoic acid requires high resolution MS1 for its unique detection with respect to 3-Oxo-1,8-octanedicarboxylic acid, demonstrated by no interference at an MS1 mass isolation window set at 25 ppm and their similar fragmentation patterns. **(B)** L-Threonine produces distinct MS2 spectra for its unique detection with regards to L-β-Homoserine, demonstrated by no interference with the use of an MS2 window at UIS2/UIS3, which cannot be accomplished solely using MS1 information. **(C)** Using the NIST 17 LC/MS library (10186 compounds) as background, UIS simulations were conducted for each analyte (9156 queries) to investigate the quantitative comparison of common acquisition methods in metabolomics (Table S-1). The following MS1/MS2 mass isolation windows were set (in daltons – Da or parts per million of 1 Da - ppm): MS1-only: 25 ppm/-; Multiple Reaction Monitoring (MRM): 0.7 Da/0.7 Da; and Data-Independent Acquisition (DIA): 25 Da/ 25 ppm, 25 ppm/25 ppm. The percentage of compounds in the NIST 17 library with no interference (the background for each query based on the given parameters), known as the percentage of unique compounds (y-axis), was calculated for each method (x-axis). *For UIS analyses, MS1-only methods used only a single precursor XIC (XIC – extracted ion chromatogram).

#### MS1 and MS2 Contribute Orthogonally to Unambiguous Detection of Complex Metabolites

We simulated four different acquisition modes: MS1, MRM, and DIA with/without a high resolution MS1 scan. For our MS1 analysis, we assumed that high resolution MS1 scans (at least 40 000 resolution or 25 ppm) would be acquired followed by extracted ion chromatogram (XIC) analyses. For our targeted metabolomics analysis, we assumed a standard QQQ instrument to be used with a 0.7 quadrupole isolation window. For DIA data we first simulated a method using a high resolution MS1 scan, followed by several high resolution fragment ion scans (at least 25 ppm) with a large (25 Da) quadrupole isolation window analyzed by both MS1 and MS2 XIC analysis. A second method was also simulated for DIA, only reliant on the high resolution MS2 scans (no MS1). Our simulations demonstrate that both accurate precursor and fragment ion information contribute orthogonally to unambiguous compound detection in complex samples (Fig. 2). By using high resolution precursor and fragment ion information, we observe improvements in the unique detection of these metabolites, maximizing the overall percentage of unique compounds detected.

First, we investigated the effects of mass accuracy on the unique detection of compounds using two representative examples: 3-Hydroxydodecanoic acid and L-Threonine. For the compound 3-Hydroxydodecanoic acid (associated with fatty acid metabolic disorders), accuracy at the MS1 level allows unique detection with respect to 3-Oxo-1,8-octanedicarboxylic acid, which has a similar fragmentation pattern but differs in precursor *m/z* by over 300 ppm (217.1798 and 217.1071 respectively). These compounds are thus resolvable with a high resolution MS1 scan (within 25 ppm) but not separable using a standard quadrupole mass filter (MS1-only), with masses of 216.1725 Da and 216.0998 Da respectively. The similarity between their MS2 spectra highlights the importance of acquiring high resolution MS spectra, as only a high resolution MS1 precursor scan can distinguish the two analytes, while even utilizing three high resolution MS2 fragments will not uniquely map to a single analyte in the background library (Fig. 2A). In our second example, L-Threonine and L-β-Homoserine produce distinct MS2 spectra that are easily distinguishable using the second most abundant fragment ion filtering even with a low resolution QQQ instrument (comparing high-quality transitions greater than 10% of the maximum relative intensity in the fragment spectrum), but cannot be distinguished using only MS1 information, as these amino acids have the same precursor *m/z* (mass of 119.0582 Da, precursor *m/z* of 120.0655) but differ in their fragmentation patterns. For this pair of metabolites, their unique fragment ion spectra specifically provides optimal discriminating power, as methods using MS2 data show no interference (Fig. 2B). These examples highlight the combined importance of both high resolution precursor and high resolution fragment ion *m/z* to provide high selectivity for analyte detection.

#### DIA Outperforms MRM and MS1-only Metabolomics Methods with respect to Unambiguous Detection

Next, we compared our four simulated analytical methods on the full NIST 17 library by using each of the 9156 compounds (filtered for common adducts) as a query against the full set of 10186 compounds (21334 unique precursors) in the library. When quantitatively comparing different metabolomics methods, our simulations show that DIA data acquisition followed by the extraction of ion chromatograms with narrow mass tolerances (25 ppm MS1 XIC / 25 ppm MS2 XIC) outperformed both MS1-only and MRM-based methods with respect to unambiguous detection, reducing the number of ambiguous compounds by ~2.8-fold and ~1.8-fold respectively (76.4% ambiguous compounds for MS1-only, 49.2% for MRM UIS_3_, and 27.4% for DIA with MS1 UIS_3_, Fig. 2C). Our analysis demonstrates that neither MS1-based extraction at 25 ppm accuracy nor a single transition in MRM (0.7 Da MS1/0.7 Da MS2) is sufficient to uniquely detect most of the compounds in the NIST 17 library. This demonstrates that neither reliance on few transitions (as in MRM) nor reliance on MS1 signal alone is sufficient for unambiguous compound detection using mass alone. Interestingly, MRM assays performed comparably to DIA when using fragment ion information only. While extracting a single fragment ion only performed worse than using accurate precursor information both in MRM and DIA, the selectivity of fragment-ion based analysis can be boosted substantially by simply adding additional qualifier ions (e.g. selecting multiple fragment ions) for analysis (see difference between UIS_1_ and UIS_3_ in Fig. 2). Interestingly, we find that MS2-based DIA analysis performs highly similar to MRM both measured when using a single transition (83.6% vs. 82.5% non-unique compounds) or using three transitions (50.9% vs. 49.2%). However, in DIA analysis (but not MRM), specificity can be further enhanced using accurate precursor information, which then outperforms both MRM and MS1 (rightmost bars in Fig. 2C). Overall, we show that both MRM and DIA outperform MS1-only analysis in terms of specificity, while DIA enhanced with high-resolution MS1 data outperforms both MS1-only and MRM-based analyses, due to its capability of extracting both high resolution precursor and fragment ion traces for any analyte of interest.

#### Theoretical Saturation of Compound Uniqueness based on Number of Transitions, Matrix Complexity and Collision Energy

In order to estimate the relative contribution of individual factors to ambiguous detection in metabolomics, we performed simulations where we varied the transition number, background complexity and collision energy. Using our UIS framework, we were able to quantify the effects of the number of transitions, matrix complexity, and collision energy on unambiguous detection.

Experimentally, additional selectivity can often be achieved by increasing the number of monitored transitions (qualifier ions). However, this may result in a higher limit of detection and lower throughput (for MRM), thus increasing cost. To evaluate this tradeoff quantitatively, we assessed the theoretical saturation of unique compounds based on the number of transitions utilized for UIS measures (Fig. 3A). While DIA methods (with accurate MS1 information) are close to saturation already when using the 3 most abundant transitions and only small gains can be achieved by using more than 3 fragment ion transitions, this was not the case for methods without a high resolution MS1 scan (traditional targeted metabolomics approaches on a QQQ instrument and DIA without accurate MS1 information). The relative difference between MRM and DIA (with MS1) was 42.0% when using 1 transition and 16.2% when using up to 8 transitions, indicating that using more transitions in MRM closes the gap between MRM and DIA. However even when using 8 transitions, MRM failed to uniquely detect 36.0% of analytes (compared to the 19.8% of ambiguous analytes when using DIA with the precursor ion trace). Since about half of the NIST library compounds have less than 5 high quality transitions (Fig. 3B), the use of minimal transitions available is important to determine methods that perform best with limited data, a common occurrence in clinical literature. Our findings indicate that specifically for compounds with relatively few fragment ions, DIA strongly outperforms traditional QQQ platforms in terms of assay selectivity, highlighting the potential of this method in untargeted clinical studies (Fig. 3B).

**Figure 3.**
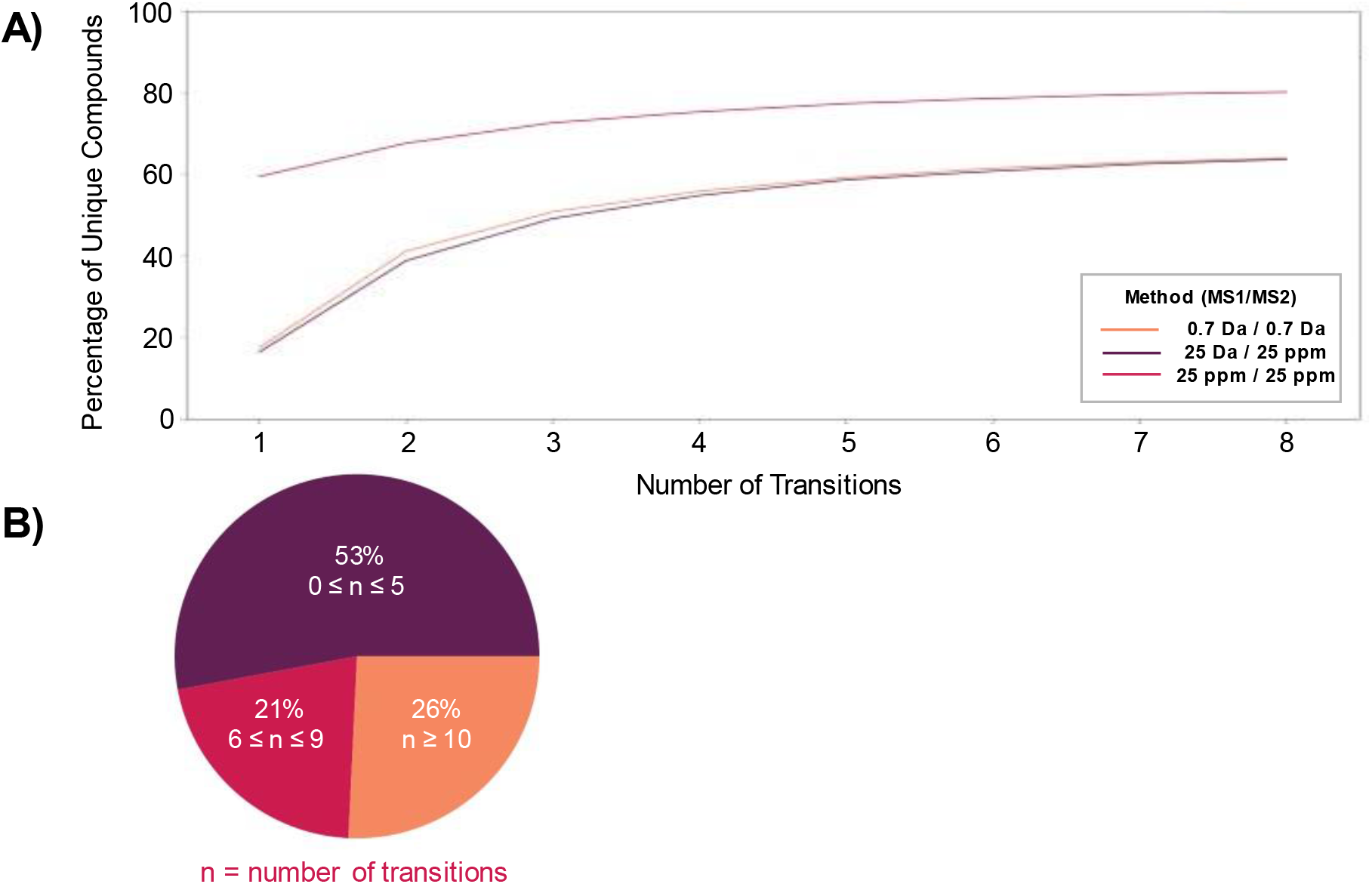
Theoretical Saturation of Unique Compounds based on the Number of Transitions. UIS simulations were performed for different acquisition methods using MS2 information (MRM, DIA), in relation to the number of transitions (*n)* defined for UISn ranging from *n*=1-8 (x-axis). Mass isolation windows were set (MS1/MS2) in daltons (Da) or parts per million (ppm) of 1 Da (Table S-1). **(A)** A saturation effect is observed at 36.0% non-uniquely detected analytes for MRM (0.7 Da/0.7 Da), 36.3% for DIA without MS1 (25 Da/25 ppm), and 19.8% for DIA with MS1 (25 ppm/25 ppm). Using more than three transitions maximizes the percentage of unique compounds for DIA with MS1, while saturation starts at more than five transitions for MRM and DIA without MS1 (<3% difference in ambiguous detection). **(B)** The total number of transitions for each spectrum in the NIST 17 LC/MS library is displayed (post-filtering of the background metabolome with transitions greater than 10% of the maximum relative intensity in fragment spectra). About 50% of the background library contains ≤ 5 transitions, in which D has reduced assay redundancy in comparison to MRM with minimal transitions available.

Next, we investigated whether our findings were dependent on the specific sample matrix (~9000 compounds) chosen and whether our results would change in samples of different background complexity or by restricting the number of considered compounds. Restricting analysis to a particular subset of compounds is equivalent to the practically employed approach of restricting compound detection to a narrow window of chromatographic retention time, thus effectively removing a large amount of potentially interfering background analytes. We simulated this by randomly choosing subsets of the NIST 17 library to produce background sample matrices of lower complexity (ranging from 1000-9000 compounds) and used extrapolation to estimate how a more complex sample matrix would behave (Fig. 4). We found that the relative performance of the individual methods using MS2 (MRM using a QQQ and DIA) is independent of the sample complexity, demonstrated by the level of saturation of each method in regards to the percentage of uniquely detected compounds. This demonstrates that our findings are not dependent on the compound library size, but are likely to be generalizable for a wide range of analytical and sample conditions (with corresponding increased sample complexity), and would likely also hold when analysis is restricted to a small region of the chromatography. Comparably, MS1-profiling methods demonstrate a higher dependence on the background composition, with saturation at larger sample complexities. This demonstrates the importance of additional separation methods (like RT) for MS1-only approaches. Analyzing increasing sample complexity, we found saturation beginning at around ~8,000 compounds in the background for MS1-profiling methods and around ~3,000-5,000 compounds for MS2-based methods (defined by <3% difference), resulting in 87.0% non-uniquely detected analytes for MS1-only, 54.8% for MRM, and 30.3% for DIA when including the precursor ion trace at UIS_3_. These observed saturation behaviors may indicate that our results would not change even for more complex backgrounds, assuming that the structural composition of more complex samples are comparable to the one studied here.

**Figure 4.**
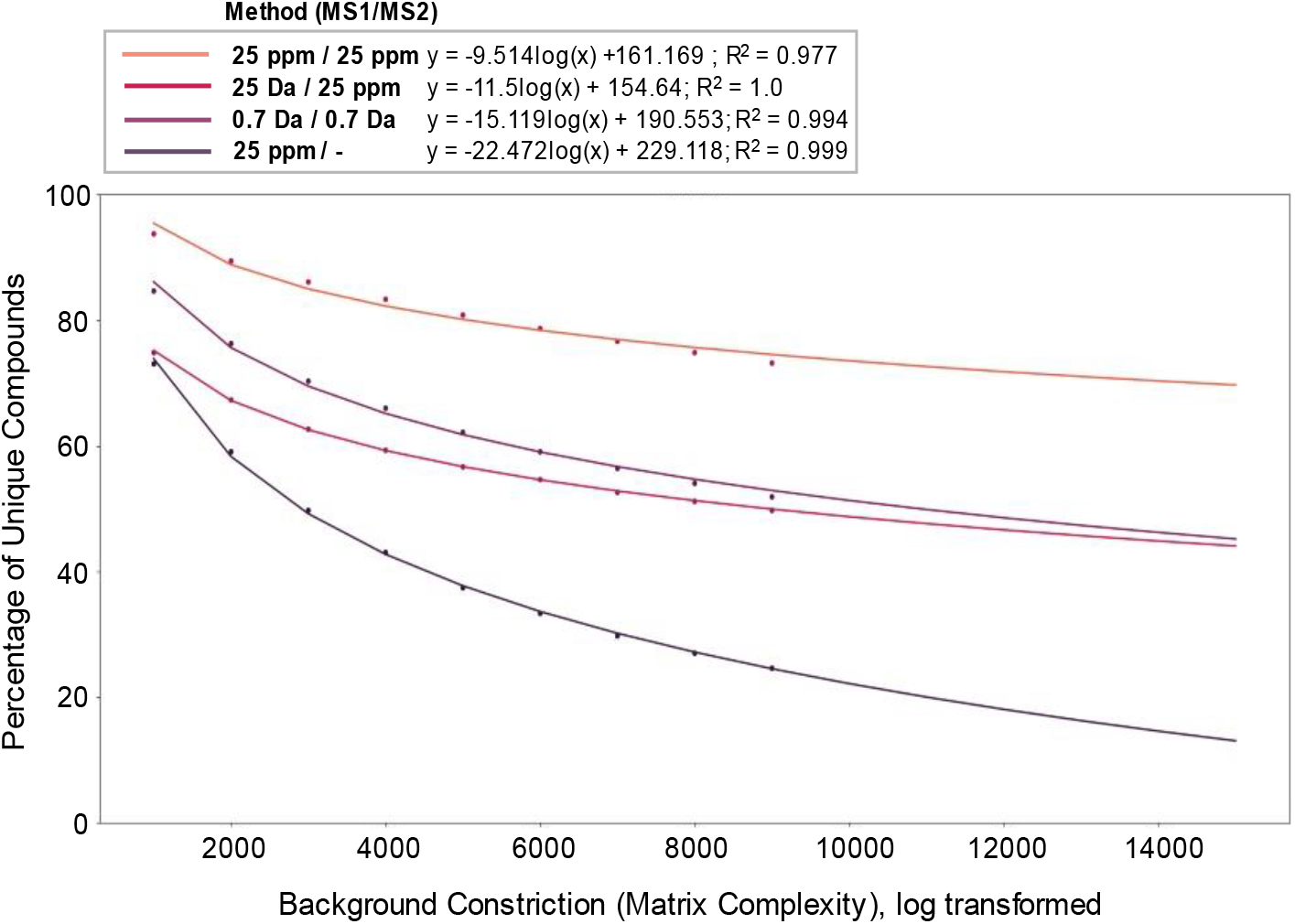
Theoretical Saturation of Unique Compounds based on Sample Matrix Complexity. Simulations were conducted to determine the performance of different MS methods (measured by the percentage of unique compounds, y-axis), in relation to different matrix complexities (x-axis). Mass isolation windows were set (MS1/MS2) in daltons (Da) or parts per million (ppm) of 1 Da (Table S-1). For methods using MS2, three transitions were used (UIS3). Simulations were performed for 100 samples of each subset from the NIST 17 LC/MS library as background (varying from 1000-9000 compounds), with extrapolation to 15,000 compounds. A saturation effect is observed at 87.0% non-uniquely detected analytes for MS1-only (25 ppm/-), 54.8% for MRM (0.7 Da/0.7 Da), 55.9% for DIA without MS1 (25 Da/25 ppm), and 30.3% for DIA with MS1 (25 ppm/25 ppm).

Finally, we studied the effect of collision energy (CE) for the optimal differentiation of interfering compounds based on their fragment spectra. While our previous analyses were restricted to spectra with CE 35eV+/−5eV only, we now performed a pairwise evaluation of spectral similarity across all available collision energies in the analyte library (for Q-TOF instruments), to answer the question whether specific collision energies are particularly efficient at producing pairwise dissimilar spectra and thus allowing compound differentiation. The use of different collision energies has been previously shown to facilitate structural elucidation, findings suggesting the exploration of different collision energy settings for broader coverage of the metabolome.^18–20^ For this analysis we used a total set of 2240 compounds which had fragment ion spectra acquired with up to 29 different collision energy settings, resulting in ~3,000,000 spectral pairs for analysis when using an MS1-only acquisition at 25 Da (Fig. 5). For each pair of interfering compounds, we computed the collision energies at which we achieved maximally different spectra. From the compounds with interference (2234/2240 compounds), we observed that metabolites varied in the number of unique pairwise-optimal collision energies (POCE) from 1-26, and in the number of required POCE to differentiate between its interfering compounds from 1-16, highlighting the diversity of analyte pairs being compared (Fig. 5B). POCE varies substantially between analyte pairs, indicating that different collision energies provide unique information, and while one collision energy may be optimal to differentiate a target compound from another compound A, a different collision energy may be needed to differentiate it from a second compound B. Thus, our analysis suggests that choosing a single CE value for each compound would not be optimal for the analysis of diverse metabolites which differ in bond strength and have wide mass dependencies (and where normalized collision energies are commonly used to compensate for this aspect). The observed spread of required POCE per compound demonstrates the importance of measuring compounds at multiple collision energies due to the distinctive information present at different CE that aids in a compound’s unique detection hus, the observed optimal discriminating power of collision energy between interfering compounds from our analysis indicates the importance of prior CE optimization in assay design, specific to instrument type and background sample matrix.

**Figure 5.**
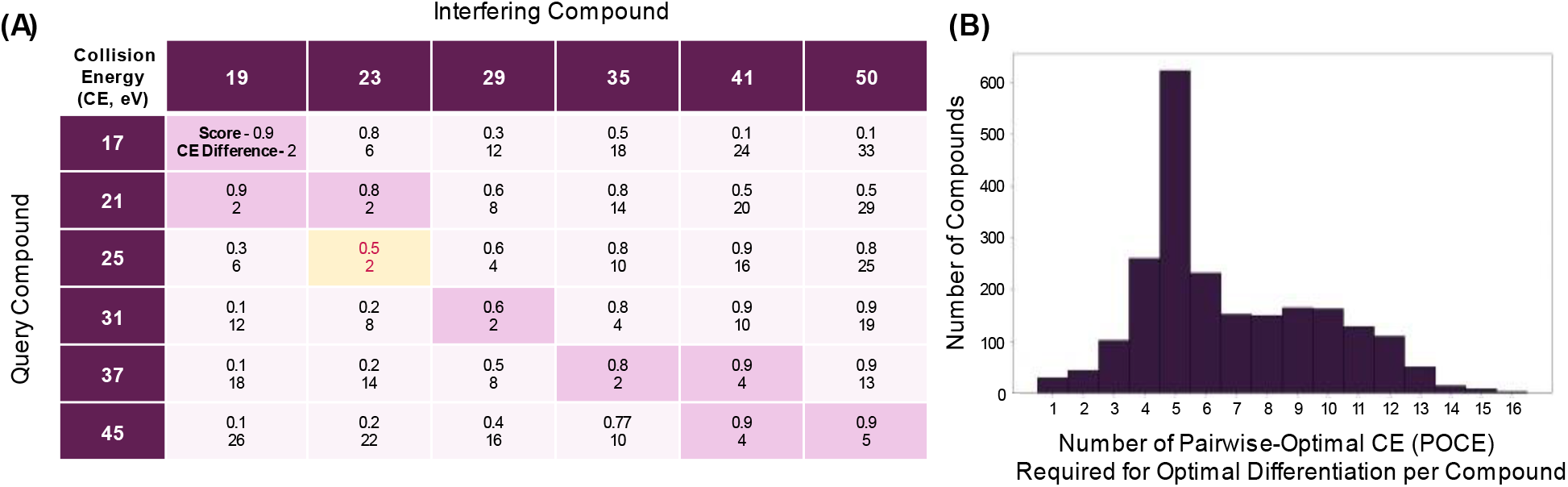
Identifying Pairwise-Optimal Collision Energies using Cosine Similarity. Collision energies (CE) were studied through pairwise evaluation of spectral similarity to determine if different CE facilitate the differentiation of interfering compounds. Compounds from the NIST 17 library measured using a Q-TOF instrument were filtered (by experimental conditions, H+ adduct, and an MS1-only mass isolation window set at 25 Da) to calculate pairwise-optimal CE for each compound. Spectra of the remaining 2234 compounds (2234/2240 had interference) and their interfering compounds acquired at varying collision energies (up to 29 different CE settings) were compared by their negative absolute difference in CE and their cosine similarity score. **A)** For example, a query compound and its interfering compound were compared and their similarity matrix is shown. Comparisons with a minimum absolute difference in CE were first chosen for each row and column (below a maximum CE difference threshold, Fig. S-8, highlighted in purple), from which a pairwise optimal CE (POCE) was selected based on the overall minimum similarity score (highlighted in yellow). **B)** The distribution of the number of POCE groups required for optimal differentiation per compound (2234 compounds) is demonstrated.

When accounting for factors of transition number and background complexity in regards to unambiguous detection, the overall theoretical saturation of a compound’s uniqueness was observed at 87.0% non-uniquely detected analytes for MS1-only, 40.0% for MRM, and 21.9% for DIA. From the analyzed methods, we include MS1-only analysis as a baseline comparison, and focus on highlighting the need for MS2 information in metabolomics analyses. One important limitation of our study is that we assume that all analytes are separable by retention time (RT) and we do not simulate patterns of exactly co-eluting fragment ions created by more than one analyte. Secondly, additional information is often used for confirmation of compound detection such as accurate retention times (derived from internal or external standards), isotopic intensity patterns or relative fragment ion intensity. This means that our values for selectivity are lower bounds and this extra information will improve selectivity across all studied acquisition methods equally, and thus not impact our overall conclusions in regards to the relative performance of the individual methods; in fact, our subsampled simulations (Fig. 4) clearly indicate that if the number of eligible interfering analytes are restricted (for example by restricting the analysis to a narrow RT window), our conclusions still hold. Similarly, in practical applications sample matrices will vary and assuming a sample matrix composed of all NIST 17 library compounds may either be too optimistic or too pessimistic for a particular application. This includes the poor coverage of lipids in the NIST 17 dataset that may lack representation of common lipid fragmentation patterns amongst isomers (i.e. double bonds, modifications and total carbon number), as well as the presence of bioactive peptides; these compounds should ideally be treated separately from other metabolites. However, while the sample matrix will influence the absolute number of unambiguously detectable analytes, it will not impact the relative performance profile of the individual methods, which we have also shown (Fig. 4). Finally, we have not analyzed sensitivity in our analysis, where a clear tradeoff is present between sensitivity and selectivity. Although opting for wider peak isolation windows increases sensitivity in traditional data-dependent methods, here we focus on the study of targeted methods where small isolation window sizes are more common. Specifically in this study, high resolution precursor isolation windows (25 ppm) have been studied in addition to 0.7 Da windows, representative of traditional isolation behaviour in triple quadrupole instruments.^21^

We demonstrate that an acquisition method using both precursor and fragment ion XIC at 25 ppm accuracy (DIA) is sufficient to unambiguously detect a large number of structurally heterogeneous analytes in a complex sample matrix of over 10,000 compounds. This work uses simulations to showcase DIA as an emerging MS acquisition method with high selectivity for metabolomics, consistent with previous experimental work in the field.^5,8,20,25–26^ Here we quantitatively show the impact that orthogonal information contributed from both MS1 and MS2 levels have on compound detection and differentiation from background compounds using *m/z* separation. Additionally, these findings support the need for an efficient pipeline for DIA analysis, building upon current tools in the field (i.e. MetaboDIA, MetDIA, MS-DIAL).^22–24^ Since our method only requires *m/z* coordinates as input, our conclusions are independent of lab-specific factors and chromatographic setup. We further demonstrate that the relative performance of the studied acquisition methods is consistent over a wide number of parameters, such as sample complexity and chromatographic constriction of the search space. In addition to the criterion of *m/z*, we expect additional orthogonal sources of information to maximize unique detection in the metabolome. This is supported by previous studies demonstrating the benefits of utilizing additional information for detection, such as retention time, collision cross-section and isotopic abundance patterns (Fig. S-4).^27^

#### Conclusions

We use unique ion signature analyses to study the performance characteristics of several widely used acquisition methods in metabolomics, and demonstrate the benefit of using both high resolution precursor and high resolution fragment ion *m/z* for unambiguous compound detection on a set of over 10,000 compounds. Our study highlights the potential of DIA for unambiguous compound detection (and quantification) in complex samples. Overall, we provide a global perspective on unambiguous compound detection and present a robust framework to study this phenomenon quantitatively.

## Acknowledgements

This project was funded by the Government of Canada through CIHR (FRN 166094) as well as NSERC (CREATE grant #528163-2019). P.S and H.R. conceived the project. All authors supplied ideas to the experimental design. P.S. performed the data analysis and prepared the publication. A.Y. performed additional analyses and helped prepare the publication.

H.R. supervised the project. All authors discussed the results and edited the paper, with additional thanks to Olga Zaslaver and Oliver Alka for further discussion and edits. H.R. is supported by the Canadian Foundation for Innovation and is the Canada Research Chair in Mass Spectrometry-based Personalized Medicine.

## Supporting Information

**Table S-1.** Simulation of Mass Isolation at MS1 and MS2 levels. **Table S-2.** Analysis of Interfering Compounds for Representative Examples at MS1 and MS2 Levels. **Table S-3.** UIS Results – Theoretical Saturation for MS1 and MS2 levels at 1 ppm accuracy. **Table S-4.** UIS Results - Influence of HCD vs. CID on UIS. **Figure S-1.** UIS Simulations of Common Acquisition Methods in Metabolomics. **Figure S-2.** Number of Transitions per Query (UIS_3_) in NIST 17. **Figure S-3.** NIST 17 Background Metabolome *m/z* Distribution. **Figure S-4.** Theoretical Saturation for MS1 and MS2 levels at 1 ppm accuracy. **Figure S-5.** UIS Comparison without Query MS2 Maximum Relative Intensity Filter. **Figure S-6.** Influence of HCD vs. CID on UIS. **Figure S-7.** NIST 17 Background Metabolome Distribution by Instrument Type and Collision Energy. **Figure S-8.** Defining Pairwise-Optimal Collision Energies. **Figure S-9.** Distribution of the Number of POCE required for Optimal Differentiation of Compounds Measured using Q-TOF vs. HCD instruments. **Figure S-10.** Validation of UIS Pipeline using NIST 20. **Figure S-11.** Theoretical Saturation of Unique Compounds in NIST 20. **Figure S-12.** Unique Ion Signature Simulations. **Text S-1.** The UIS Concept: Metabolomics vs. Proteomics.

