## Supplementary Information for "Analyzing Assay Specificity in Metabolomics using Unique Ion Signature Simulations"

| Method | MS1 Mass Isolation Window Size | MS2 Mass Isolation Window Size |
| --- | --- | --- |
| MS1-only (QQQ) | 0.7 Da | - |
| MS1-only (high resolution) | 25 ppm | - |
| MRM | 0.7 Da | 0.7 Da |
| PRM | 2 Da | 20 ppm |
| DIA (SWATH) | 25 Da | 25 ppm |
| DIA (SWATH with high resolution MS1) | 25 ppm | 25 ppm |

**Table S-1. Simulation of Mass Isolation at MS1 and MS2 levels.** Using the NIST 17 LC/MS library (10186 compounds) as background, simulations were conducted for each analyte (9156 queries) by setting different mass tolerances associated with different MS methods. The following MS1/MS2 mass isolation windows were set (in daltons – Da or parts per million of 1 Da - ppm): MS1-only: 0.7 Da/- (QQQ), 25 ppm/- (high resolution); Multiple Reaction Monitoring (MRM): 0.7 Da/0.7 Da; Parallel Reaction Monitoring (PRM): 2 Da/20 ppm; and Data-Independent Acquisition (DIA - SWATH): 25 Da/25 ppm (no MS1 data), 25 ppm/25 ppm (high resolution MS1 extraction).

| Query: 3-Hydroxydodecanoic acid | | | |
|---|---|---|---|
| MS2-level Interference (at $UIS_3$) | | | |
| *Compound Name* | *Precursor m/z* | *Adduct* | *Structure* |
| (-)-Isolongifolol | 205.1951 | $[M+H-H_20]+$ | 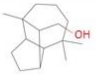 |
| α-Bisabolol | 205.1951 | $[M+H-H_20]+$ | 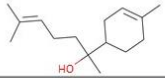 |
| DL-α-Lipoamide | 206.0668 | [M+H]+ | 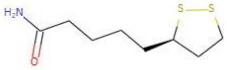 |
| Dodecanedioic acid | 213.1485 | $[M+H-H_20]+$ | 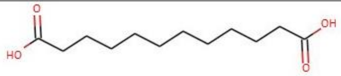 |
| 3-Oxo-1,8-octanedicarboxylic acid | 217.1071 | [M+H]+ | 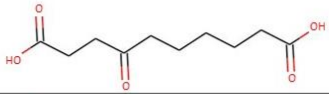 |
| Undecanedioic acid | 217.1434 | [M+H]+ | 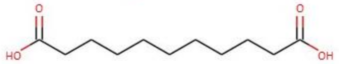 |
| Sebacic acid monomethyl ester | 217.1434 | [M+H]+ | 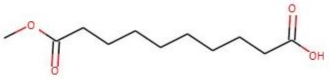 |
| Query: L-Threonine | | | |
| MS1-level Interference | | | |
| *Compound Name* | *Precursor m/z* | *Adduct* | *Structure* |
| O-Methyl-DL-serine | 120.0655 | [M+H]+ | 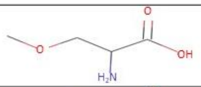 |
| L-β-Homoserine | 120.0655 | [M+H]+ | 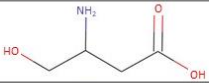 |
| N-Methyl-L-serine | 120.0655 | [M+H]+ | 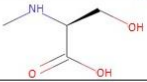 |
| O-t-Butyl-L-serine, methyl ester | 120.0655 | $[M+H-C_4H_8]+$ | 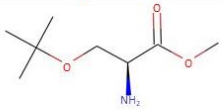 |

**Table S-2. Analysis of Interfering Compounds for Representative Examples at MS1 and MS2 Levels**. Pairs of queries and their corresponding interfering compounds at the MS1 and MS2 levels are shown (Fig. 2A-B). 3-Hydroxydodecanoic acid requires high resolution MS1 for its unique detection, demonstrated by the listed MS2-level interference at $UIS_3$ (0.7 Da/0.7 Da, 25 Da/25 ppm). L-Threonine produces distinct MS2 spectra for its unique detection, with interference at the MS1 level (25 ppm/-). Interfering compounds in the NIST 17 library can be analyzed for any compound of choice using scripts found at https://github.com/premyshan/DIAColliderMetabo.

| Saturation at MS1 Level | | | | | | | |
|---|---|---|---|---|---|---|---|
| MS1 Mass Isolation Window Size | MS2 Mass Isolation Window Size | Percentage of Unique Compounds | | | Percentage of Ambiguous Compounds | | |
| 25 ppm | - | 23.56 | | | 76.44 | | |
| 10 ppm | - | 34.10 | | | 65.90 | | |
| 1 ppm | - | 41.48 | | | 58.52 | | |
| **Saturation at MS2 Level** | | | | | | | |
| MS1 Mass Isolation Window Size | MS2 Mass Isolation Window Size | Percentage of Unique Compounds | | | Percentage of Ambiguous Compounds | | |
| | | $UIS_1$ | $UIS_2$ | $UIS_3$ | $UIS_1$ | $UIS_2$ | $UIS_3$ |
| 25 Da | 25 ppm | 16.38 | 38.81 | 49.15 | 83.62 | 61.19 | 50.85 |
| 25 Da | 10 ppm | 20.14 | 41.32 | 51.62 | 79.86 | 58.68 | 48.38 |
| 25 Da | 1 ppm | 40.08 | 65.03 | 74.20 | 59.92 | 34.97 | 25.80 |
| 25 ppm | 25 ppm | 59.47 | 67.62 | 72.63 | 40.53 | 32.38 | 27.37 |
| 25 ppm | 10 ppm | 61.98 | 69.95 | 74.73 | 38.02 | 30.05 | 25.27 |
| 25 ppm | 1 ppm | 79.01 | 86.13 | 90.17 | 20.99 | 13.87 | 9.83 |
| 10 ppm | 10 ppm | 62.59 | 70.33 | 75.04 | 37.41 | 29.67 | 24.96 |
| 1 ppm | 1 ppm | 79.43 | 86.31 | 90.33 | 20.57 | 13.69 | 9.67 |

**Table S-3. UIS Results - Theoretical Saturation for MS1 and MS2 levels at 1 ppm accuracy.** Using the NIST 17 LC/MS library (10186 compounds) as background, simulations were conducted for each analyte (9156 queries) by setting restrictive mass tolerances (up to 1 ppm) at the MS1 and/or MS2 level (Fig. S-4). The percentage of unique compounds (with no interference) and the percentage of ambiguous compounds (non-uniquely identified analytes) in the NIST 17 library were calculated for each method, as shown.

| Higher Energy Dissociation (HCD) | | | | | | | |
|---|---|---|---|---|---|---|---|
| MS1 Mass Isolation Window Size | MS2 Mass Isolation Window Size | Percentage of Unique Compounds | | | Percentage of Ambiguous Compounds | | |
| 0.7 Da | - | 0.38 | | | 99.62 | | |
| 25 ppm | - | 30.16 | | | 69.84 | | |
| | | $UIS_1$ | $UIS_2$ | $UIS_3$ | $UIS_1$ | $UIS_2$ | $UIS_3$ |
| 0.7 Da | 0.7 Da | 23.03 | 49.18 | 58.13 | 76.97 | 50.82 | 41.87 |
| 25 Da | 25 ppm | 20.46 | 44.53 | 54.04 | 79.54 | 55.47 | 45.96 |
| 25 ppm | 25 ppm | 65.55 | 72.90 | 76.27 | 34.45 | 27.10 | 23.73 |

| Collision-Induced Dissociation (CID, Ion Trap Fourier Transform) | | | | | | | |
|---|---|---|---|---|---|---|---|
| MS1 Mass Isolation Window Size | MS2 Mass Isolation Window Size | Percentage of Unique Compounds | | | Percentage of Ambiguous Compounds | | |
| 0.7 Da | - | 0.33 | | | 99.67 | | |
| 25 ppm | - | 28.84 | | | 71.16 | | |
| | | $UIS_1$ | $UIS_2$ | $UIS_3$ | $UIS_1$ | $UIS_2$ | $UIS_3$ |
| 0.7 Da | 0.7 Da | 29.03 | 52.39 | 59.28 | 70.97 | 47.61 | 40.72 |
| 25 Da | 25 ppm | 34.75 | 59.82 | 65.16 | 65.25 | 40.18 | 34.84 |
| 25 ppm | 25 ppm | 66.87 | 73.91 | 77.17 | 33.13 | 26.09 | 22.83 |

**Table S-4. UIS Results - Influence of HCD vs. CID on UIS.** Using the NIST 17 LC/MS library, overlapping compounds acquired using either higher energy dissociation (HCD) or collision-induced dissociation (CID, Ion Trap Fourier Transform - IT-FT) for Orbitrap instruments were compared to investigate the influence of HCD vs. CID on unique detection using common acquisition methods (MS1-only, MRM, DIA). Simulations were conducted for each analyte (6110 queries). The percentage of unique compounds (with no interference) and the percentage of ambiguous compounds (non-uniquely identified analytes) were calculated for each method, as shown. *For UIS analyses, MS1-only methods used only a single precursor XIC (XIC – extracted ion chromatogram).

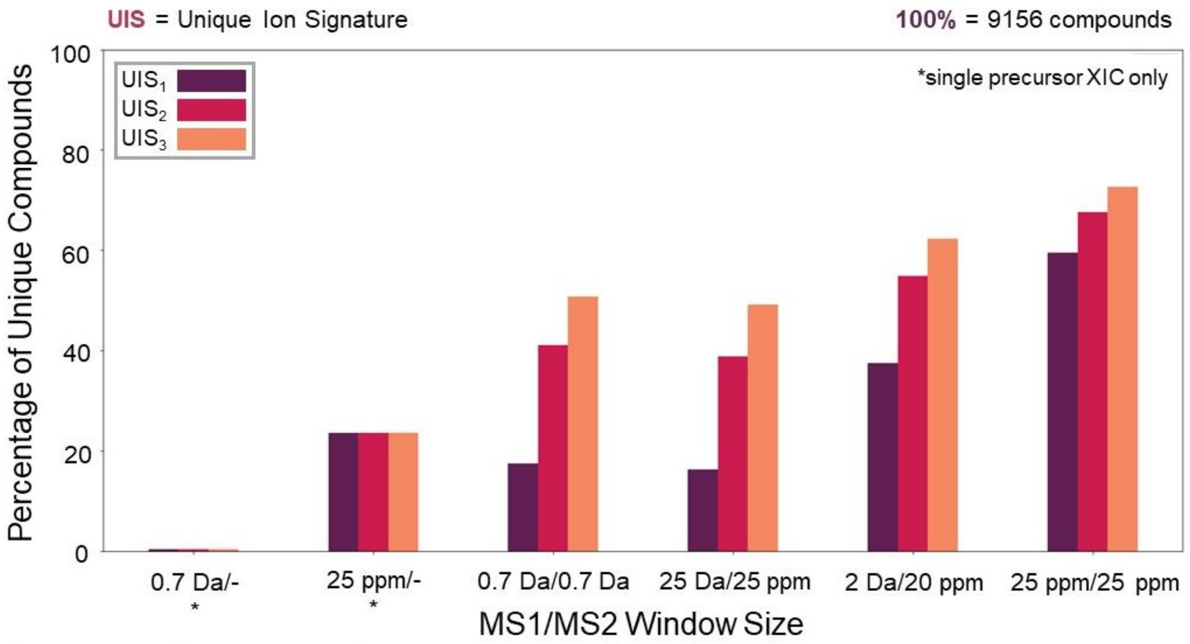

| MS1 Mass Isolation Window Size | MS2 Mass Isolation Window Size | Percentage of Unique Compounds | | | Percentage of Ambiguous Compounds | | |
|---|---|---|---|---|---|---|---|
| 0.7 Da | - | 0.49 | | | 99.51 | | |
| 25 ppm | - | 23.56 | | | 76.44 | | |
| | | $UIS_1$ | $UIS_2$ | $UIS_3$ | $UIS_1$ | $UIS_2$ | $UIS_3$ |
| 0.7 Da | 0.7 Da | 17.49 | 41.12 | 50.81 | 82.51 | 58.88 | 49.19 |
| 25 Da | 25 ppm | 16.38 | 38.81 | 49.15 | 83.62 | 61.19 | 50.85 |
| 2 Da | 20 ppm | 37.55 | 54.94 | 62.36 | 62.45 | 45.06 | 37.64 |
| 25 ppm | 25 ppm | 59.47 | 67.62 | 72.63 | 40.53 | 32.38 | 27.37 |

**Figure S-1. UIS Simulations of Common Acquisition Methods in Metabolomics.** Using the NIST 17 LC/MS library (10186 compounds) as background, simulations were conducted for each analyte (9156 queries) to investigate whether an assay can be constructed that uniquely detects the analyte within the background using common acquisition methods (MS1-only, MRM, PRM, DIA). MS1/MS2 mass isolation windows were set accordingly (in daltons - Da, or parts per million of 1 Da - ppm), as listed in Table S-1. The percentage of compounds in the NIST 17 library with no interference (the background for each query based on the given parameters), known as the percentage of unique compounds (y-axis), as well as the percentage of ambiguous compounds (non-uniquely identified analytes), were calculated for each method (x-axis). *For UIS analyses, MS1-only methods used only a single precursor XIC (XIC – extracted ion chromatogram).

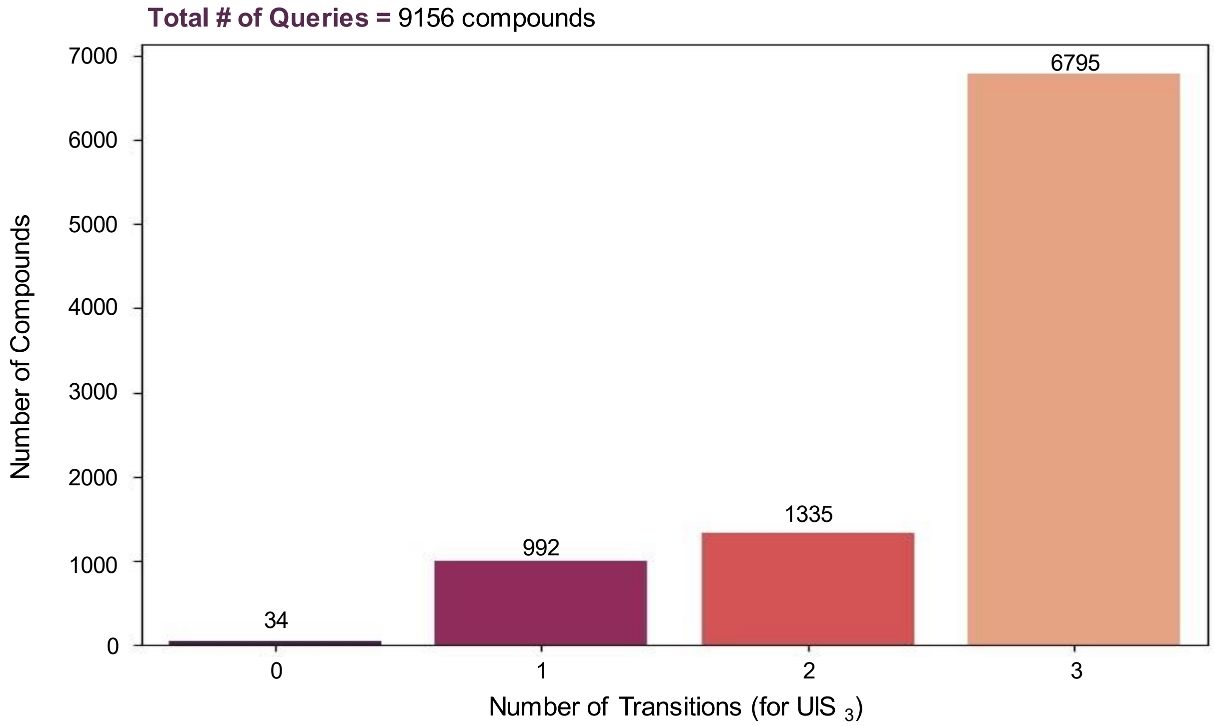

**Figure S-2. Number of Transitions per Query (UIS3) in NIST 17.** The number of transitions (x-axis) per query for a UIS3 assay is demonstrated using the NIST 17 LC/MS library (9156 queries, number of compounds on the y-axis) after filtering for data acquisition (removal of stereoisomers, positive ion mode, HCD and Q-TOF instrument type, CE=35eV +/- 5eV, H+ and Na+ adducts).

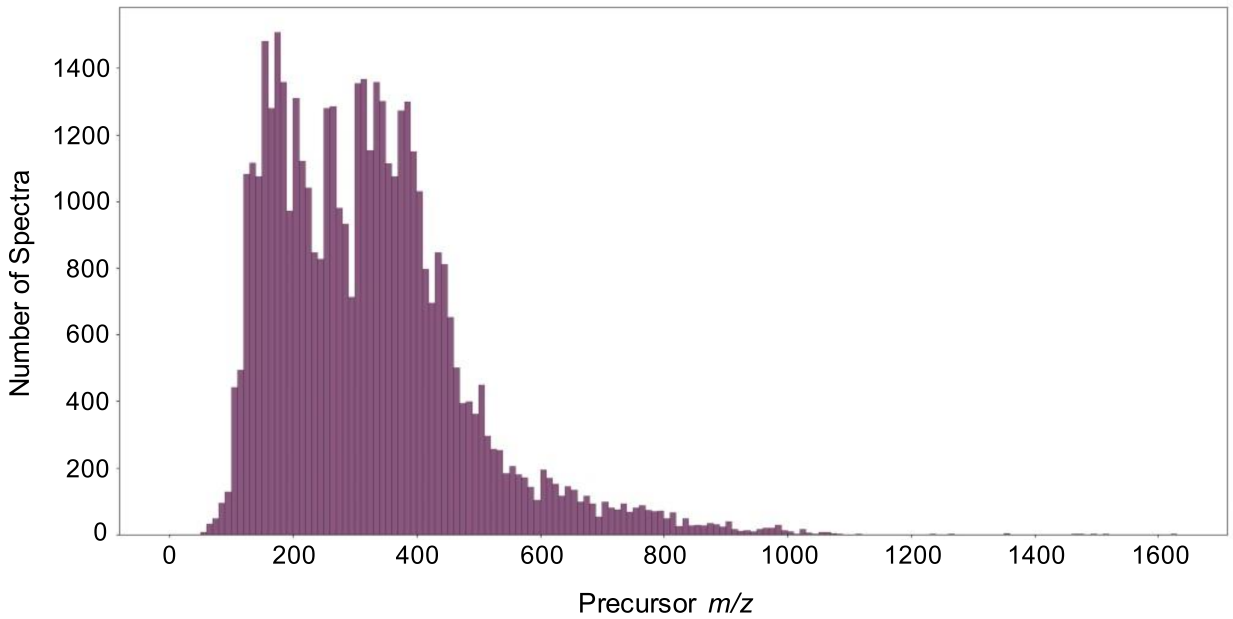

**Figure S-3. NIST 17 Background Metabolome *m/z* Distribution.** The *m/z* distribution (x-axis) of 10186 background metabolites (in the given NIST 17 metabolome used for UIS simulations) is displayed after filtering for data acquisition (removal of stereoisomers, positive ion mode, HCD and Q-TOF instrument type, CE=35eV +/- 5eV, all adducts).

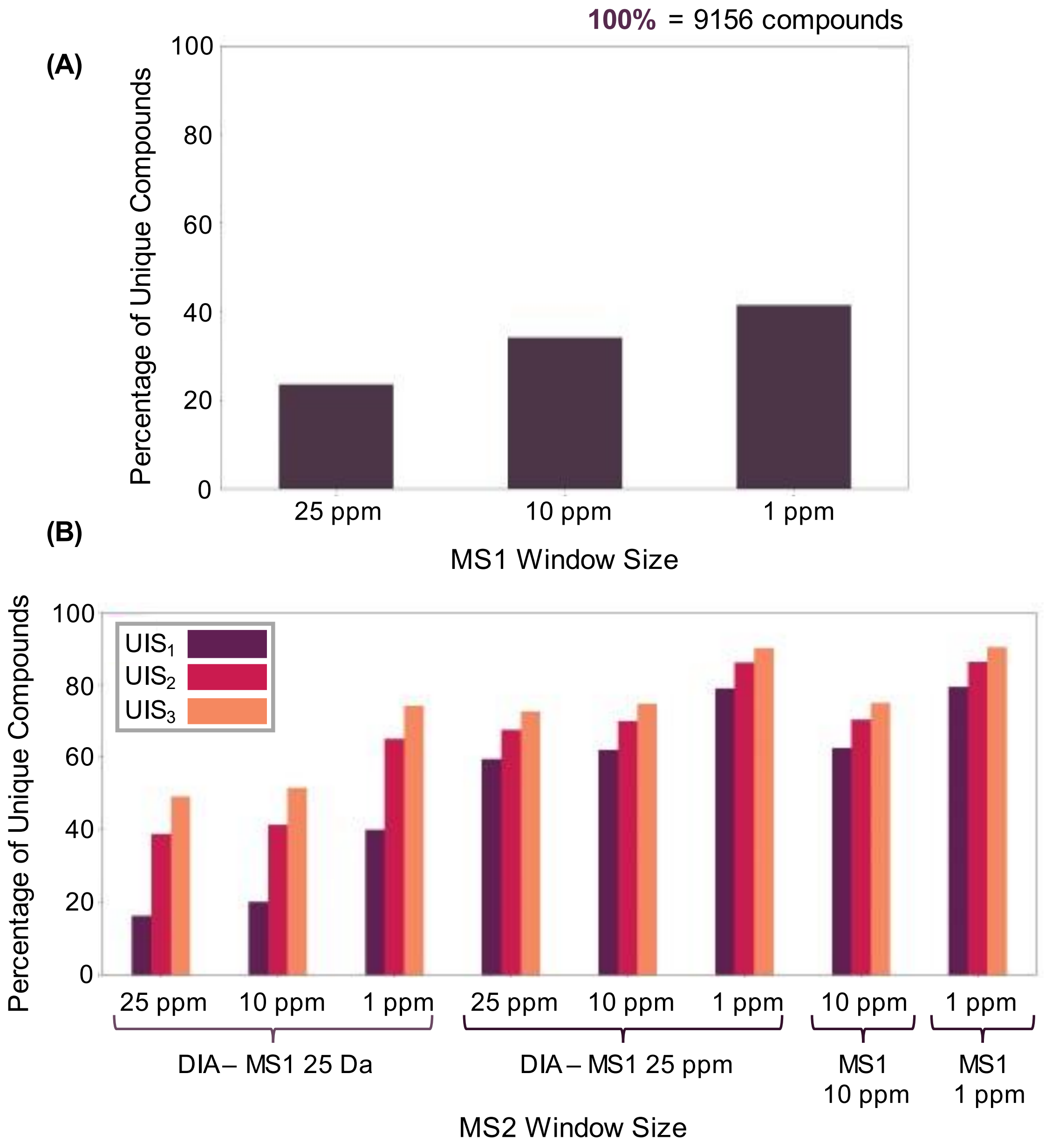

**Figure S-4. Theoretical Saturation for MS1 and MS2 levels at 1 ppm accuracy.** Using the NIST 17 LC/MS library (10186 compounds) as background, simulations were conducted for each analyte (9156 queries) with high mass accuracy by setting restrictive mass tolerances (up to 1 ppm) at the MS1 or MS2 level. **(A)** At the MS1 level, different mass tolerances were set in parts per million of 1 Da – 25 ppm, 10 ppm and 1 ppm (23.6%, 34.1% and 41.5% uniquely detected analytes respectively). **(B)** At the MS2 level, DIA with (MS1 at 25ppm) and without high resolution MS1 (MS1 at 25 Da) were simulated with the following MS2 mass isolation window settings: 25 ppm, 10 ppm and 1 ppm. Additional settings were tested (MS1/MS2) to determine saturation at both MS1 and MS2 levels: 10 ppm/10 ppm, 1 ppm/1 ppm. The percentage of compounds in the NIST 17 library with no interference (the background for each query based on the given parameters), known as the percentage of unique compounds (y-axis), was calculated for each method (x-axis), with results shown in Table S-3.

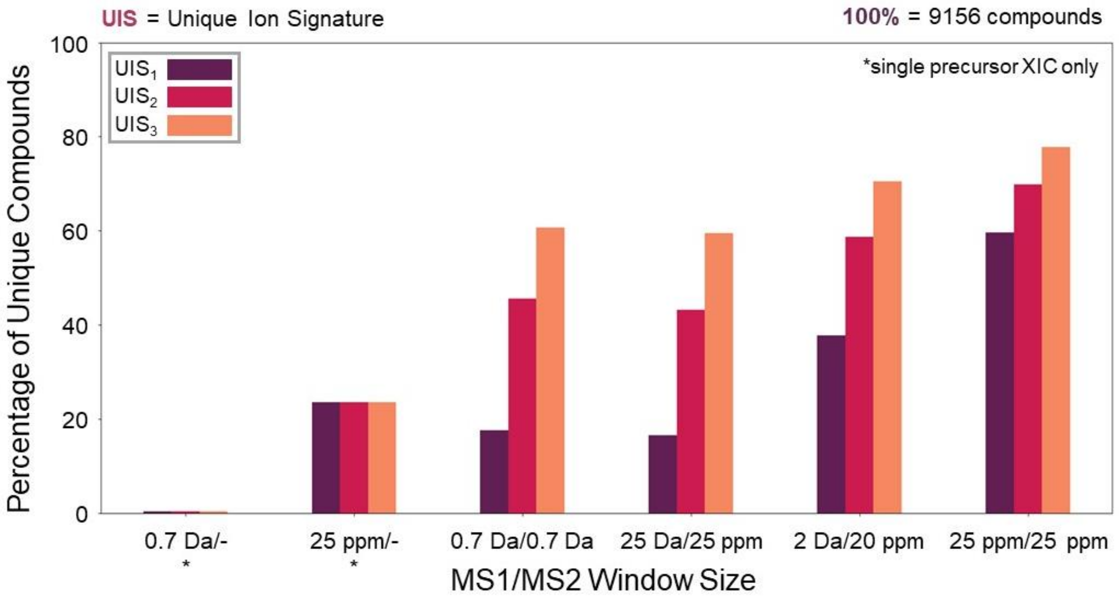

| MS1 Mass Isolation Window Size | MS2 Mass Isolation Window Size | Percentage of Unique Compounds | | | Percentage of Ambiguous Compounds | | |
|---|---|---|---|---|---|---|---|
| 0.7 Da | - | 0.49 | | | 99.51 | | |
| 25 ppm | - | 23.56 | | | 76.44 | | |
| | | $UIS_1$ | $UIS_2$ | $UIS_3$ | $UIS_1$ | $UIS_2$ | $UIS_3$ |
| 0.7 Da | 0.7 Da | 17.67 | 45.61 | 60.68 | 82.33 | 54.39 | 39.32 |
| 25 Da | 25 ppm | 16.61 | 43.17 | 59.51 | 83.39 | 56.83 | 40.49 |
| 2 Da | 20 ppm | 37.85 | 58.70 | 70.51 | 62.15 | 41.30 | 29.49 |
| 25 ppm | 25 ppm | 59.63 | 69.83 | 77.85 | 40.37 | 30.17 | 22.15 |

**Figure S-5. UIS Comparison without Query MS2 Maximum Relative Intensity Filter.** Using the NIST 17 LC/MS library (10186 compounds) as background, simulations were conducted for each analyte (9156 queries) by setting different mass tolerances associated with different MS methods (MS1-only, MRM, PRM, DIA). Fragment spectra for each query included all transitions (with no MS2 maximum relative intensity filter), allowing the use of UIS with no threshold. MS1/MS2 mass isolation windows were set accordingly (in daltons - Da, or parts per million of 1 Da - ppm), as listed in Table S-1. The percentage of compounds in the NIST library with no interference (the background for each query based on the given parameters), known as the percentage of unique compounds (y-axis), as well as the percentage of ambiguous compounds (non-uniquely identified analytes), were calculated for each method (x-axis). *For UIS analyses, MS1-only methods used only a single precursor XIC (XIC – extracted ion chromatogram).

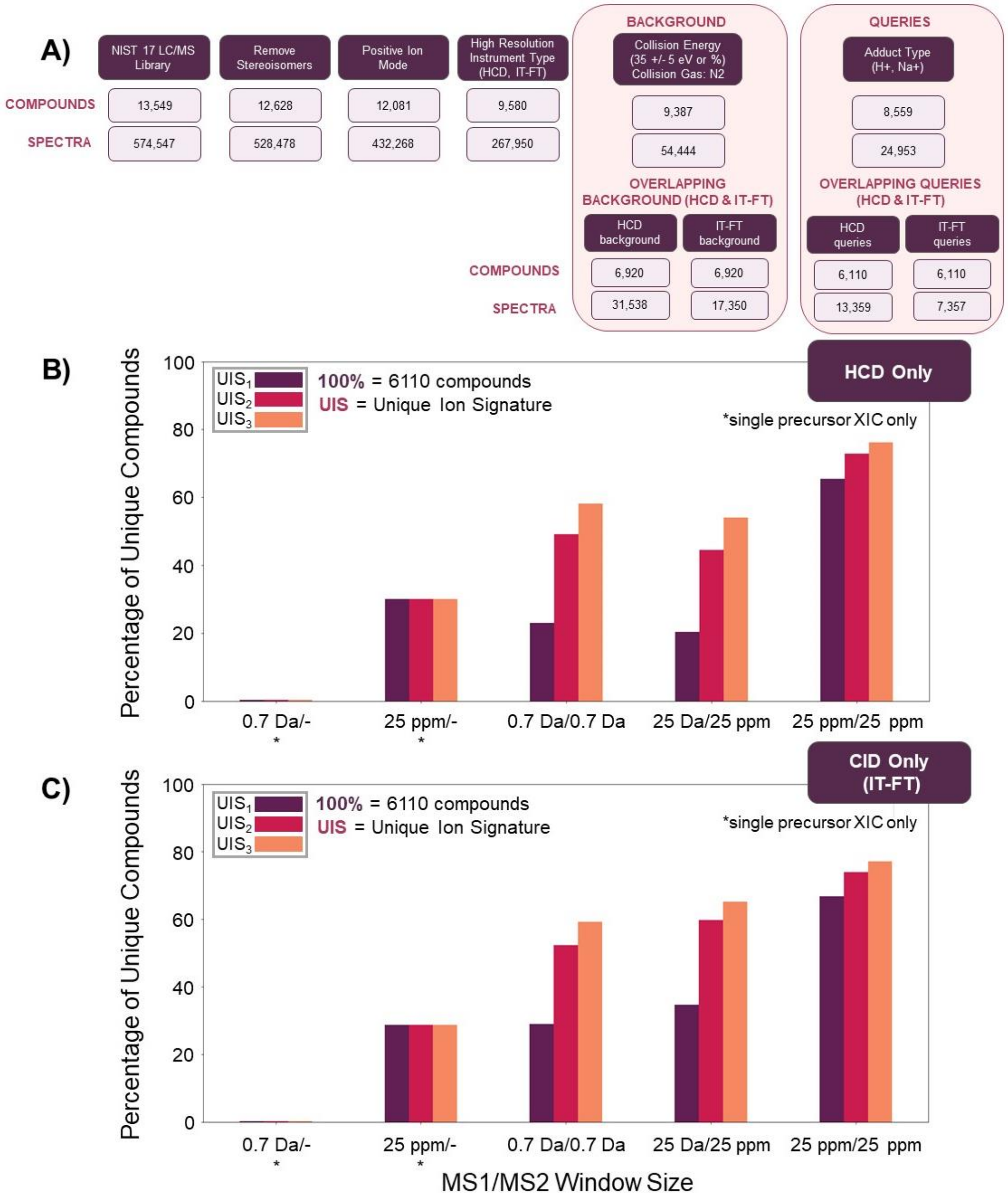

**Figure S-6. Influence of HCD vs. CID on UIS.** Using the NIST 17 LC/MS library, **(A)** overlapping compounds acquired using **(B)** higher energy dissociation (HCD) or **(C)** collision-induced dissociation (CID, Ion Trap Fourier Transform - IT-FT) for Orbitrap instruments were compared to investigate the influence of HCD vs. CID on unique detection using common acquisition methods (MS1-only, MRM, DIA). The following MS1/MS2 windows were set (in daltons - Da, or parts per million of 1 Da - ppm): MS1-only: 0.7 Da/-, 25 ppm/-; Multiple Reaction Monitoring (MRM): 0.7 Da/0.7 Da; Data-Independent Acquisition (DIA): 25 Da/ 25 ppm, 25 ppm/25 ppm. Simulations were conducted for each analyte (6110 queries) and the percentage of compounds with no interference (the background for each query based on the given parameters), known as the percentage of unique compounds (y-axis), was calculated for each method (x-axis), with results shown in Table S-4. *For UIS analyses, MS1-only methods used only a single precursor XIC (XIC – extracted ion chromatogram).

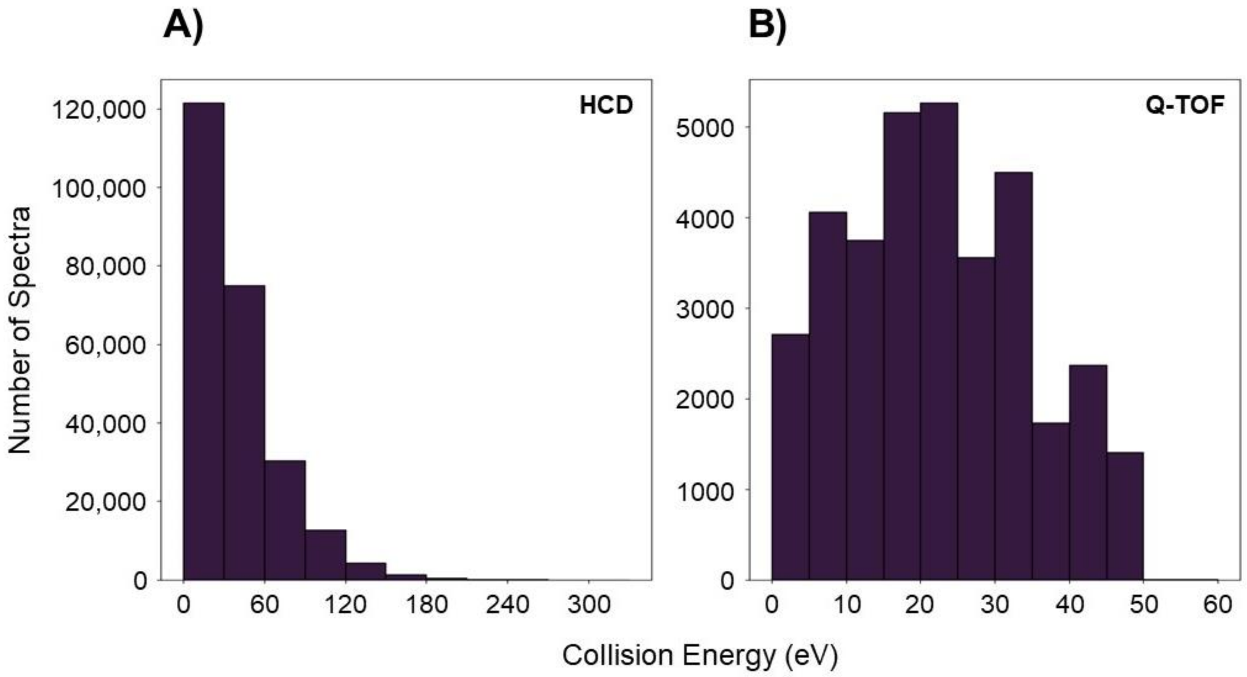

**Figure S-7. NIST 17 Background Metabolome Distribution by Instrument Type and Collision Energy.** Spectra in the NIST 17 LC/MS library (y-axis) are displayed after filtering (removal of stereoisomers, positive ion mode, high resolution instrument type). This characterization is based on instrument type - **(A)** higher energy dissociation/HCD with 284 CE settings and **(B)** quadrupole time-of-flight/Q-TOF with 29 CE settings, further specified by the distribution of acquired collision energies (measured in eV), as indicated on the x-axis.

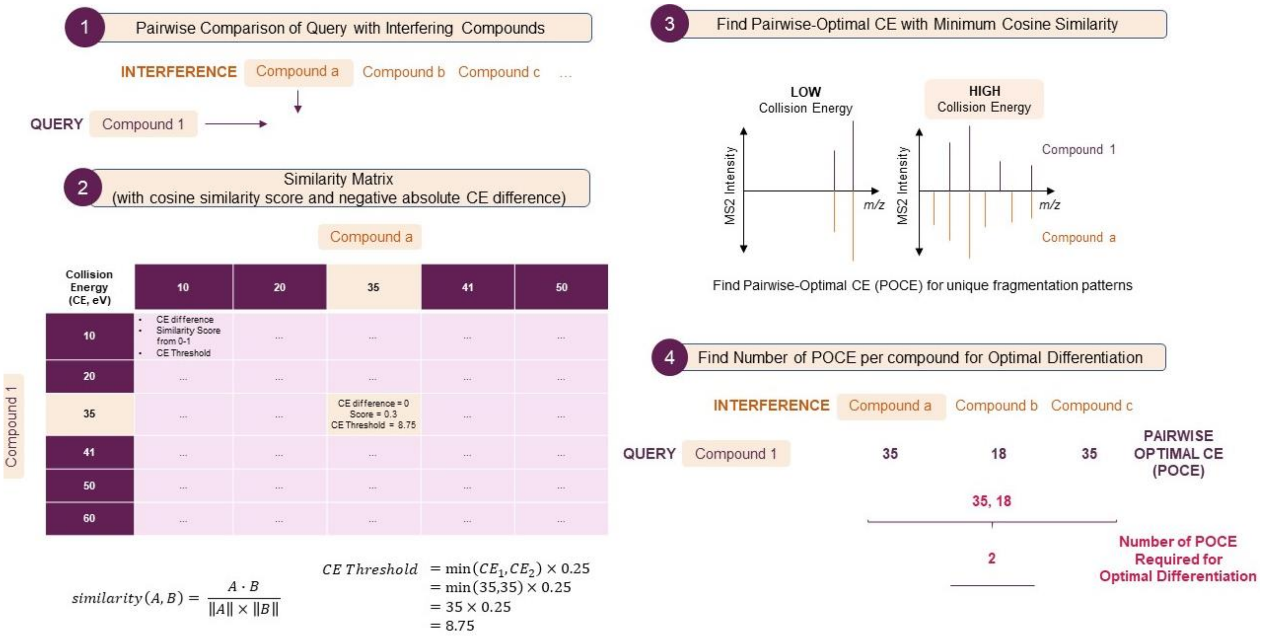

**Figure S-8. Defining Pairwise-Optimal Collision Energies.** Compounds from the NIST 17 LC/MS library were used to calculate pairwise-optimal collision energies for each metabolite. After filtering, **(1)** individual compounds were compared against each of their interfering compounds **(2)** using two values to create a similarity matrix - the negative absolute difference in CE compounds were measured at and their cosine similarity score based on their fragment spectra. First, comparisons with a minimum absolute difference in CE were chosen for each row and column. A threshold was maintained for CE difference, only keeping comparisons with an absolute CE difference less than or equal to a maximum CE difference threshold (25% of the minimum of two compared CE). **(3)** From these comparisons, a pairwise-optimal CE (POCE) was determined based on the overall minimum similarity score (highlighted in yellow). **(4)** The number of POCE required for optimal differentiation from interference was recorded for each compound of interest.

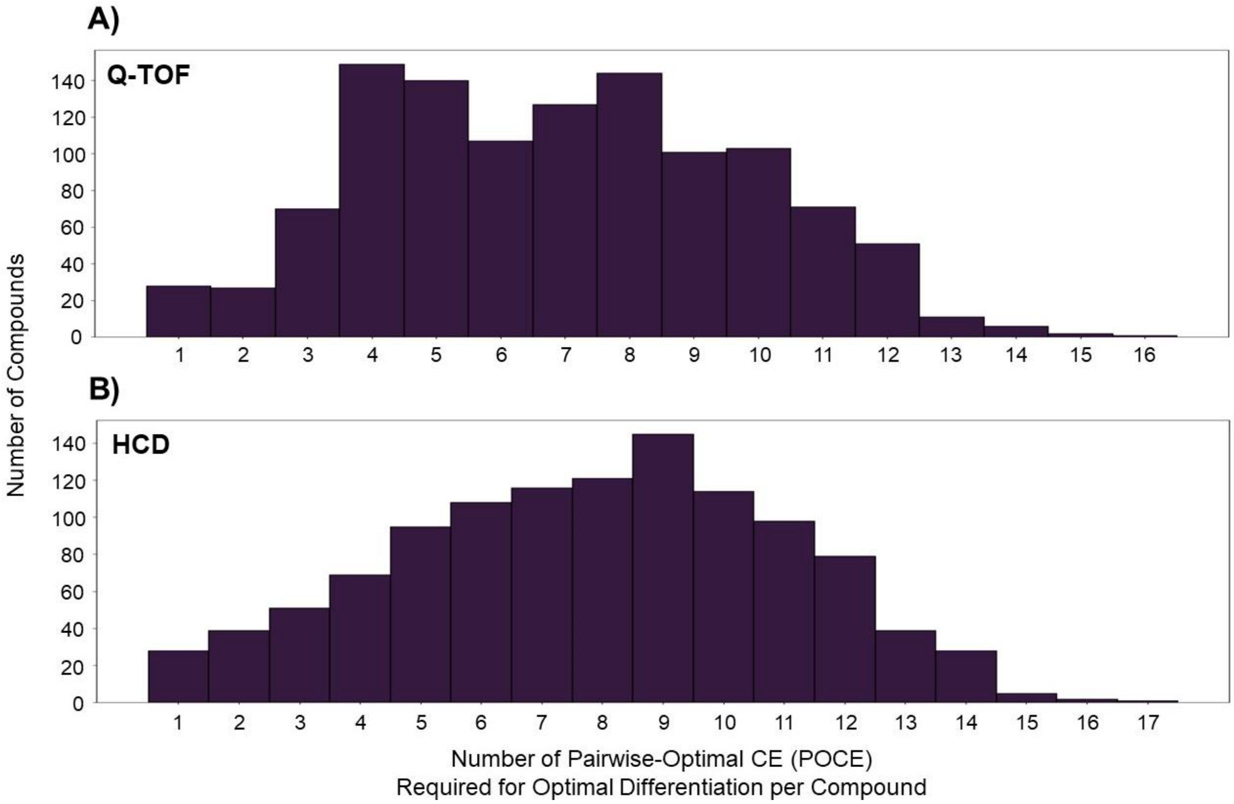

**Figure S-9. Distribution of the Number of POCE required for Optimal Differentiation of Compounds Measured using Q-TOF vs. HCD instruments.** Overlapping compounds from the NIST 17 LC/MS library (filtered for experimental conditions and H+ adduct) measured using **(A)** Q-TOF (quadrupole time-of-flight) and **(B)** HCD (higher energy dissociation) instruments were used to calculate the number of pairwise-optimal collision energies (POCE) required for optimal differentiation of each metabolite against their individual background metabolomes (1142 compounds, 1138 had interference at an MS1-only mass isolation window set at 25 Da). The distribution of the number of required POCE groups per compound is shown.

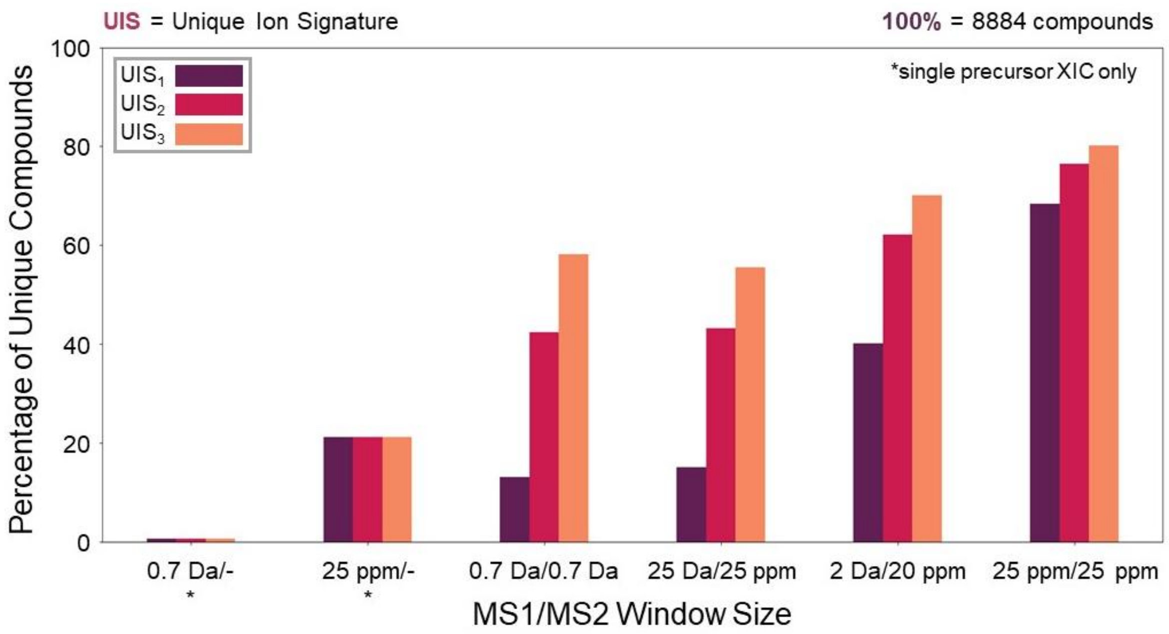

| MS1 Mass Isolation Window Size | MS2 Mass Isolation Window Size | Percentage of Unique Compounds | | | Percentage of Ambiguous Compounds | | |
|---|---|---|---|---|---|---|---|
| 0.7 Da | - | 0.64 | | | 99.36 | | |
| 25 ppm | - | 21.24 | | | 78.76 | | |
| | | $UIS_1$ | $UIS_2$ | $UIS_3$ | $UIS_1$ | $UIS_2$ | $UIS_3$ |
| 0.7 Da | 0.7 Da | 13.18 | 42.42 | 58.26 | 86.82 | 57.58 | 41.74 |
| 25 Da | 25 ppm | 15.08 | 43.22 | 55.59 | 84.92 | 56.78 | 44.41 |
| 2 Da | 20 ppm | 40.21 | 62.12 | 70.17 | 59.79 | 37.88 | 29.83 |
| 25 ppm | 25 ppm | 68.43 | 76.44 | 80.26 | 31.57 | 23.56 | 19.74 |

**Figure S-10. Validation of UIS Pipeline using NIST 20.** Using novel compounds in the NIST 20 LC/MS library (9697 compounds, 22421 precursors after removal of compounds already present in NIST 17) as background, simulations were conducted for each analyte (8884 queries) to validate UIS analyses performed on the NIST 17 library comparing common acquisition methods (MS1-only, MRM, PRM, DIA, Fig. S-1). MS1/MS2 mass isolation windows were set accordingly (in daltons - Da, or parts per million of 1 Da - ppm), as listed in Table S-1. The percentage of compounds with no interference (the background for each query based on the given parameters), known as the percentage of unique compounds (y-axis), as well as the percentage of ambiguous compounds (non-uniquely identified analytes), were calculated for each method (x-axis). *For UIS analyses, MS1-only methods used only a single precursor XIC (XIC – extracted ion chromatogram).

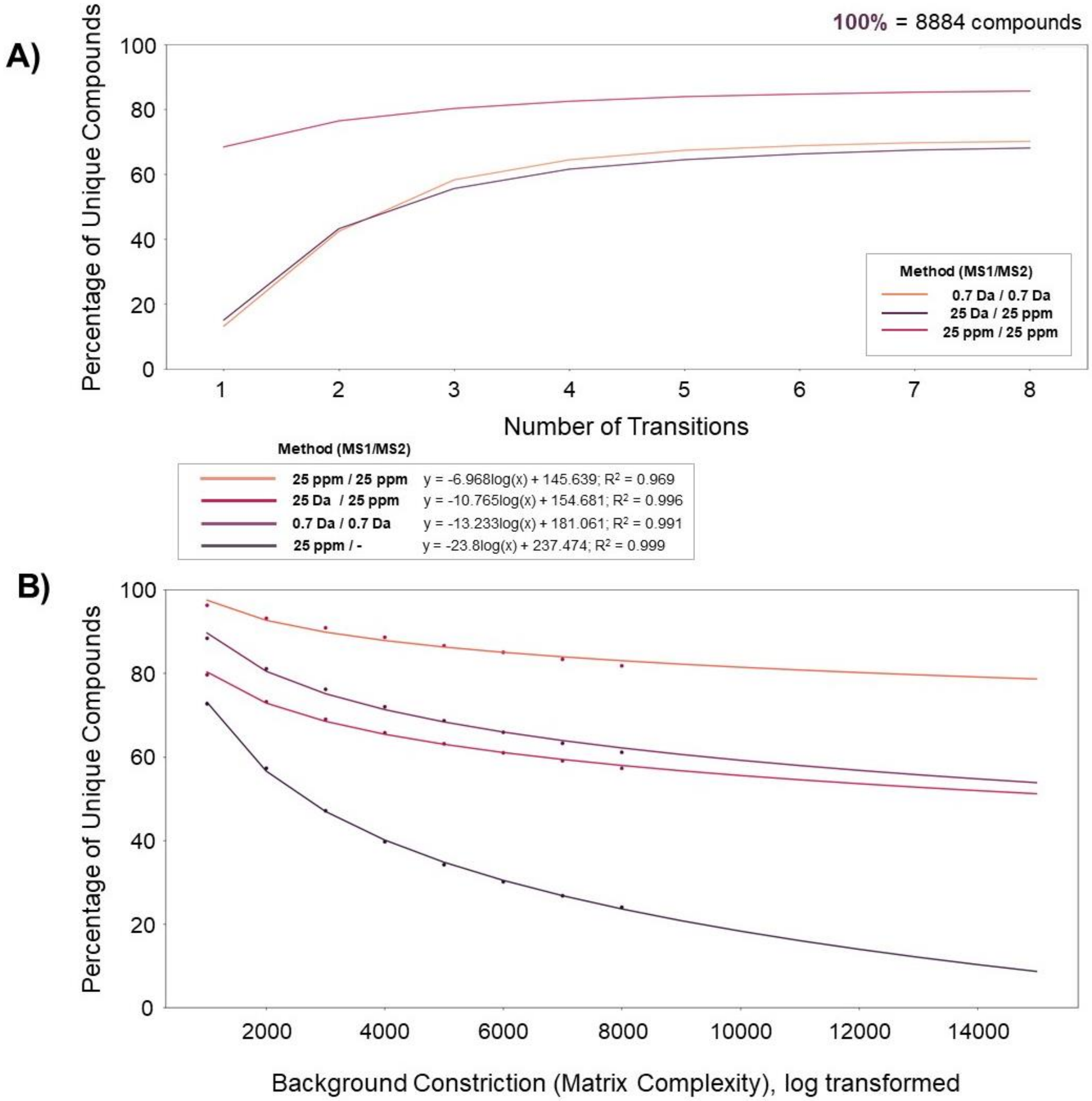

**Figure S-11. Theoretical Saturation of Unique Compounds in NIST 20.** Simulations were conducted using novel compounds in the NIST 20 LC/MS library (9697 background compounds and 8884 queries after removal of compounds already present in NIST 17) to validate UIS analyses performed on the NIST 17 library in regards to the performance of different MS methods (MS1-only, MRM, DIA, Fig. 3-4). The percentage of unique compounds (y-axis) was measured in relation to **(A)** the number of transitions (*n*) used for $UIS_n$ and **(B)** varying matrix complexities (as defined by constricting the number of compounds in the background metabolome, from 1000-8000 compounds). The following MS1/MS2 windows were set (in daltons - Da, or parts per million of 1 Da - ppm): MS1-only: 25 ppm/-; Multiple Reaction Monitoring (MRM): 0.7 Da/0.7 Da; Data-Independent Acquisition (DIA): 25 Da/ 25 ppm, 25 ppm/25 ppm (DIA with high resolution MS1). For methods using MS2, three transitions were used ($UIS_3$). For **(A),** a saturation effect is seen at 29.9% non-uniquely detected analytes for MRM, 31.9% for DIA without MS1, and 14.4% for DIA with MS1. For **(B),** a saturation effect is seen at 91.4% non-uniquely detected analytes for MS1-only, 46.2% for MRM, 48.8% for DIA without MS1, and 21.4% for DIA with MS1.

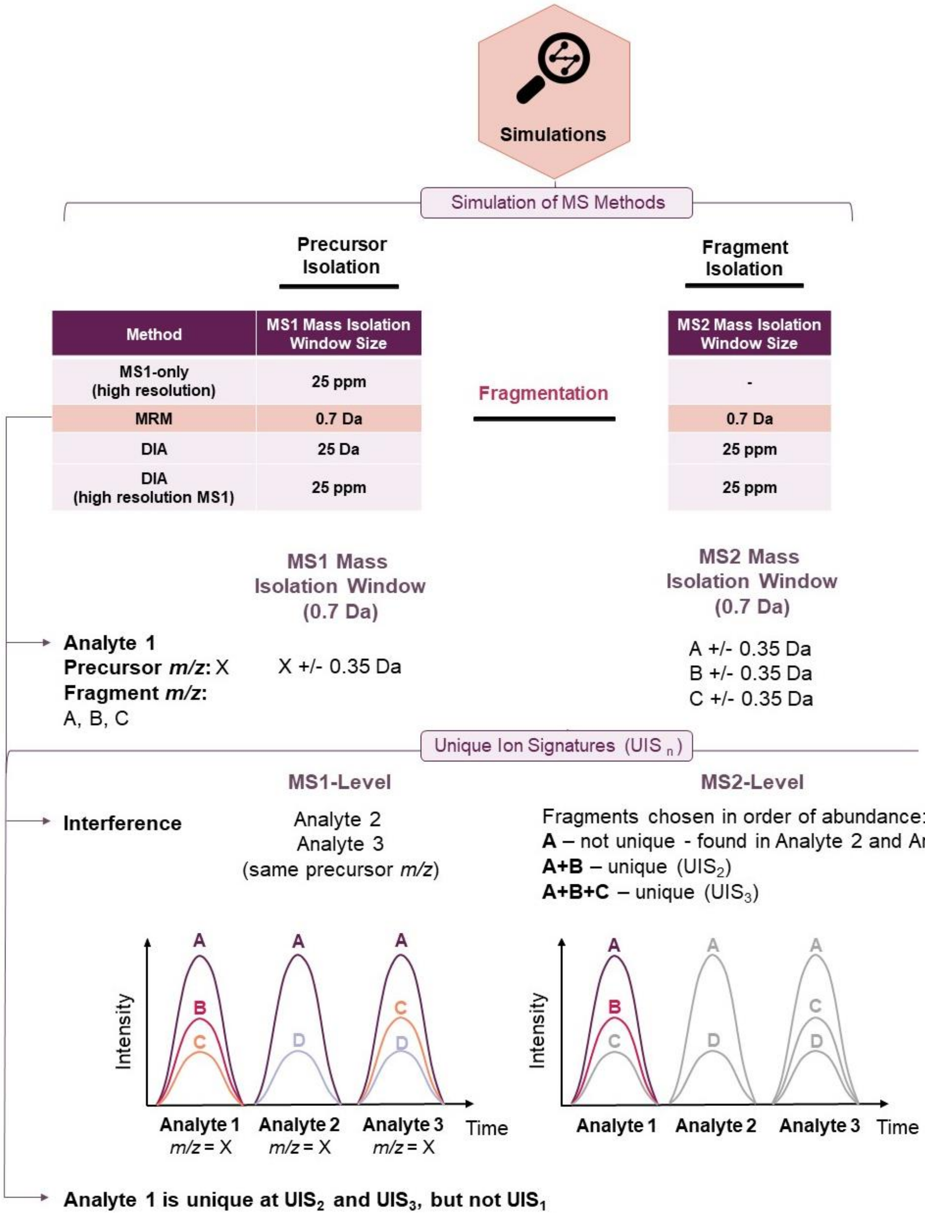

**Figure S-12. Unique Ion Signature Simulations.** Using the NIST 17 LC/MS library (9156 queries, 10186 background metabolites), unique ion signature (UIS) simulations were performed by setting realistic mass tolerances associated with different MS methods (Table S-1). For example, an MRM for Analyte 1 (with precursor *m/z* X) would contain the following three transitions: A, B, and C. Isolation window sizes were set accordingly (0.7 Da/0.7 Da for MRM), centered at the corresponding precursor and fragment *m/z* values of the query (+/- 0.35 Da), with the number of transitions defined by a value of *n* ($UIS_n$, with transitions chosen in order of abundance). Our simulations demonstrate that at the MS1-level, Analyte 1 would have the following interfering compounds: Analyte 2 and Analyte 3 (as they have the same precursor *m/z*). At the MS2-level, Analyte 1 has no $UIS_1$, a $UIS_2$ of A and B, and a $UIS_3$ of A, B, and C. Note that transition A is not a unique ion signature because this signal can be explained by either Analyte 1, Analyte 2 or Analyte 3. Figure adapted from: Röst, H.; Malmström, L.; Aerbersold, R. *Mol. Cell. Proteom*. **2012**.

**Text S-1. The UIS Concept: Metabolomics vs. Proteomics**

For our analyses, we have applied the Unique Ion Signatures (UIS) technique for the metabolomics field to address the long-standing problem of ambiguous detection not studied before to this extent. The UIS concept was initially developed to analyze uniqueness of peptide fragmentation.[14-15] In comparison to the theoretical application of this concept in the field of proteomics, we have expanded the UIS approach and adopted it for the metabolomics field (for the first time in over 12 years since its initial publication) where theoretical spectra cannot be easily predicted as in proteomics, and where collision energy optimization is an integral part of the analytical process.

Accordingly, our study is based on one of the largest collections of experimental data available: the NIST 20 library with over ~1,000,000 experimental fragment ion spectra. While a peptide can easily be compared against any other peptide through comparison of pre-computed theoretical ions (such as b and y ions), for metabolites we restricted ourselves to experimental spectral comparisons in order to compute UIS due to the poor confidence in current efforts to predict fragment ion intensities from chemical structures. Our innovation of using a large-scale database of experimental fragment ion spectra adopts the approach to metabolomics and makes it fit-for-purpose, allowing us to gain a series of novel insights.

Overall, we are introducing a novel way of comparing acquisition methods using a new concept in the field of metabolomics. Our work allows, for the first time, the comparison of targeted methods using a sound theoretical framework and shows that DIA is comparable or even superior to traditional MRM (with high resolution MS1).
